# Multiple Mechanisms Inactivate the LIN-41 RNA-Binding Protein to Ensure A Robust Oocyte-to-Embryo Transition in *Caenorhabditis elegans*

**DOI:** 10.1101/378398

**Authors:** Caroline A. Spike, Gabriela Huelgas-Morales, Tatsuya Tsukamoto, David Greenstein

## Abstract

In the nematode *Caenorhabditis elegans,* the conserved LIN-41 RNA-binding protein is a translational repressor that coordinately controls oocyte growth and meiotic maturation. LIN-41 exerts these effects, at least in part, by preventing the premature activation of the cyclin-dependent kinase CDK-1. Here we investigate the mechanism by which LIN-41 is rapidly eliminated upon the onset of meiotic maturation. Elimination of LIN-41 requires the activities of CDK-1 and multiple SCF-type ubiquitin ligase subunits, including the conserved substrate adaptor protein SEL-10/Fbw7/Cdc4, suggesting that LIN-41 is a target of ubiquitin-mediated protein degradation. Within the LIN-41 protein, two non-overlapping regions, Deg-A and Deg-B, are individually necessary for LIN-41 degradation; both contain several potential phosphodegron sequences, and at least one of these sites is required for LIN-41 degradation. Finally, Deg-A and Deg-B are sufficient, in combination, to mediate SEL-10-dependent degradation when transplanted into a different oocyte protein. Although LIN-41 is a potent inhibitor of protein translation and M-phase entry, the failure to eliminate LIN-41 from early embryos does not result in the continued translational repression of LIN-41 oocyte mRNA targets. Based on these observations, we propose a molecular model for the elimination of LIN-41 by SCF^SEL-10^ and suggest that LIN-41 is inactivated before it is degraded. Furthermore, we provide evidence that another RNA-binding protein, the GLD-1 tumor suppressor, is regulated similarly. Redundant mechanisms to extinguish translational repression by RNA-binding proteins may both control and provide robustness to irreversible developmental transitions, including meiotic maturation and the oocyte-to-embryo transition.

## INTRODUCTION

The genetic pathways controlling developmental decisions have evolved to be robust to perturbations stemming from environmental change, nutrient deprivation, and endogenous genetic variation (reviewed by Hammerstein *et al.* 2005; Félix and Wagner 2008). At a genetic level, robustness stems in large part from redundancy of control and feedback and feed-forward regulatory mechanisms built into the pathways. In turn, redundancy in genetic control also provides a hardy and permissive substrate to support evolutionary change (reviewed by Prince and Pickett 2002; Vavouri *et al.* 2008). Ultimately, the robustness of genetic pathways is central to the preservation of the germline, the immortal cell lineage required for sexual reproduction and perpetuation of a species. Cell fate decisions in the germline fall into two broad classes, those that are plastic and those that represent irreversible all-or-none commitments. For example, in several organisms the differentiated progeny of germline stem cells can dedifferentiate to repopulate the stem cell niche in response to adverse conditions that deplete the stem cell pool (Brawley and Matunis 2004; Kai and Spradling 2004; Nakagawa *et al.* 2007, 2010; Cheng *et al.* 2008; Barroca *et al.* 2009). By contrast, the commitment of an oocyte to complete meiosis and undergo fertilization represents an irrevocable decision. Here we explore the molecular genetic mechanisms controlling the commitment to fertilization during the final stages of oogenesis in the nematode *Caenorhabditis elegans.*

In *C. elegans,* as in many animals, fully grown oocytes are transcriptionally quiescent and depend on a maternal load of protein and messenger RNA to complete their development. As a consequence, the dramatic cell cycle and developmental changes that occur during the transition from oogenesis to embryogenesis are driven by post-transcriptional mechanisms. Such mechanisms include protein phosphorylation, the elimination of maternally-provided proteins or mRNAs, and the regulation of maternal mRNA translation (reviewed by Verlhac *et al.* 2010; Robertson and Lin 2015; Svoboda *et al.* 2017). The oocyte-to-embryo transition (OET) initiates when oocytes exit meiotic prophase and enter the first meiotic metaphase (MI), a cell cycle and developmental event also known as meiotic resumption or meiotic maturation. The OET completes when zygotic gene transcription begins after fertilization in the early embryo.

Pioneering studies using amphibian oocytes established that oocyte meiotic maturation is initiated by the activation of maturation-promoting factor (MPF), in response to progesterone from the follicle cells (Masui and Markert 1971; reviewed by Masui 2001). The principal components of MPF are the cyclin-dependent kinase Cdk1 catalytic subunit and a cyclin B regulatory subunit(Dunphy *et al.* 1988;Gautier *et al.* 1988, 1990; Lohka *et al.* 1988; reviewed by Nurse 1990). In *Xenopus,* which represents the best-studied system from a biochemical standpoint, MPF activation involves the translation of multiple, apparently redundantly-acting factors, including the cMos protein kinase, B-type cyclins, RINGO/Speedy, and proteins that remain to be identified (Kobayashi *et al.* 1991; Minshull *et al.* 1991; Nebreda *et al.* 1995; Frank-Vaillant *et al.* 1999; Haccard and Jessus 2006a; reviewed by Haccard and Jessus 2006b). Once activated, MPF stimulates multiple positive feedback mechanisms, including the activation of the Greatwall kinase, which phosphorylates and activates the protein phosphatase 2A (PP2A) inhibitor α-endosulfine (Yu *et al.* 2006; Zhao *et al.* 2008; Von Stetina *et al.* 2008; Castilho *et al.* 2009; Mochida *et al.* 2010). The inhibition of PP2A results in the activation of the CDC25 phosphatase, which removes the inhibitory CDK1 phosphorylations at Thr14 and Tyr15 catalyzed by the Weel or Mytl kinases (Kornbluth *et al.* 1994; Mueller *et al.* 1995). The initial signal from MPF activation is amplified in a feed-forward mechanism in which active CDK promotes the inactivation of its inhibitors, Weel and Mytl (reviewed by Ferrell 1999a,b), and stimulates its activator, CDC25 (Kumagai and Dunphy 1996). This regulatory mechanism generates the “switch-like” activation of MPF that promotes the rapid and irreversible cell cycle transition from prophase to metaphase (reviewed by O’Farrell *et al.* 2001; Kishimoto 2015).

MPF is the master regulator of cell cycle progression during oocyte meiotic maturation in *C. elegans* as in all examined species (Boxem *et al.* 1999; Burrows *et al.* 2006; van der Voet *et al.* 2009), yet MPF activation is regulated somewhat differently than in *Xenopus.* For example, although the inhibitory Weel/Mytl kinase Wee-1.3 is crucially important for inhibiting MPF activity in immature *C. elegans* oocytes (Burrows *et al.* 2006), an apparent Greatwall homolog is not found in the *C. elegans* genome and α-endosulfine/ensα-1 is not required for viability or fertility (Kim *et al.* 2012). Likewise, the signal that triggers MPF activation for meiotic maturation in *C. elegans* is not progesterone, but rather the major sperm protein (MSP), an abundant cytoskeletal protein that is released from sperm (Miller *et al.* 2001; Kosinski *et al.* 2005). The latter control mechanism, which serves to link meiotic maturation and ovulation to sperm availability, likely evolved in gonochoristic predecessors of facultative hermaphroditic nematode species like *C. elegans.* The rate of meiotic maturation declines substantially as a *C. elegans* hermaphrodite utilizes its limited supply of sperm for self-fertilization but rapidly increases upon mating (Kosinki *et al.* 2005). When sperm are absent, as in mutant hermaphrodites that do not produce sperm *(e.g., fog* mutant females), oocytes arrest for prolonged periods and the rate of production and growth of new oocytes declines until insemination (McCarter *et al.* 1999; Wolke *et al.* 2007; Govindan *et al.* 2009). This serves to preserve metabolically costly oocytes when sperm are unavailable for fertilization. Thus, the molecular mechanisms that control MPF activation must be exquisitely fine-tuned for sperm sensing.

Another commonality between the *C. elegans* and *Xenopus* systems is that MPF activation depends on translational control mechanisms, though the details differ. In *C. elegans,* large ribonucleoprotein (RNP) complexes containing the tripartite motif (TRIM)-NHL (NCL-1, HT2A, and LIN-41) RNA-binding protein LIN-41 and the tristetraprolin/TIS11-related RNA-binding proteins OMA-1 and OMA-2 (referred to collectively as the OMA proteins) are major downstream targets of MSP signaling (Spike *et al.* 2014a,b; Tsukamoto *et al.* 2017). LIN-41 is the chief determinant of the extended meiotic prophase of *C. elegans* oocytes (Spike *et al.* 2014a). In *lin-41* null mutants, pachytene-stage oocytes cellularize prematurely, activate CDK-1, aberrantly disassemble the synaptonemal complex, and enter M phase precociously, causing sterility (Matsuura *et al.* 2016; Spike *et al.* 2014a; Tocchini *et al.* 2016). CDK-1 activation causes oocytes to prematurely transcribe and express genes that are ordinarily restricted to differentiated cells and expressed after the OET (Allen *et al.* 2014; Spike *et al.* 2014a; Tocchini *et al.* 2014). LIN-41 inhibits CDK-1 activation in part through the 3’-untranslated region (UTR) mediated translational repression of the CDC-25.3 phosphatase (Spike *et al.* 2014a,b). By contrast, the OMA proteins are redundantly required for CDK-1 activation (Detwiler *et al.* 2001). In the absence of the OMA proteins, oocytes fail to undergo meiotic maturation despite the presence of sperm, resulting in sterility (Detwiler *et al.* 2001).

Genetic analysis suggests the OMA proteins promote meiotic maturation by inhibiting the function of LIN-41 in the most proximal oocyte. Two lines of molecular evidence are consistent with the idea that LIN-41 must be inactivated to promote meiotic maturation. First, LIN-41 is degraded upon the onset of meiotic maturation in response to CDK-1 activation (Spike *et al.* 2014a; Figure 1, A and B). Second, LIN-41 is a potent translational repressor, yet several of the mRNAs it associates with and represses are translated and co-expressed with LIN-41 prior to meiotic maturation in the −1 and −2 oocytes (Tsukamoto *et al.* 2017). These mRNAs include those encoding the RNA-binding protein SPN-4, which is required for development of the embryonic germline and the mesendoderm (Gomes *et al.* 2001), and MEG-1, which is a germplasm or P granule component needed for germline development (Leacock and Reinke 2008; Kapelle and Reinke 2011; Wang *et al.* 2014). By contrast, the OMA proteins are required for the translation of *spn-4* and *meg-1* transcripts in proximal oocytes, providing a molecular mechanism by which the OMA proteins might antagonize LIN-41 function (Tsukamoto *et al.* 2017).

**FIGURE 1.**
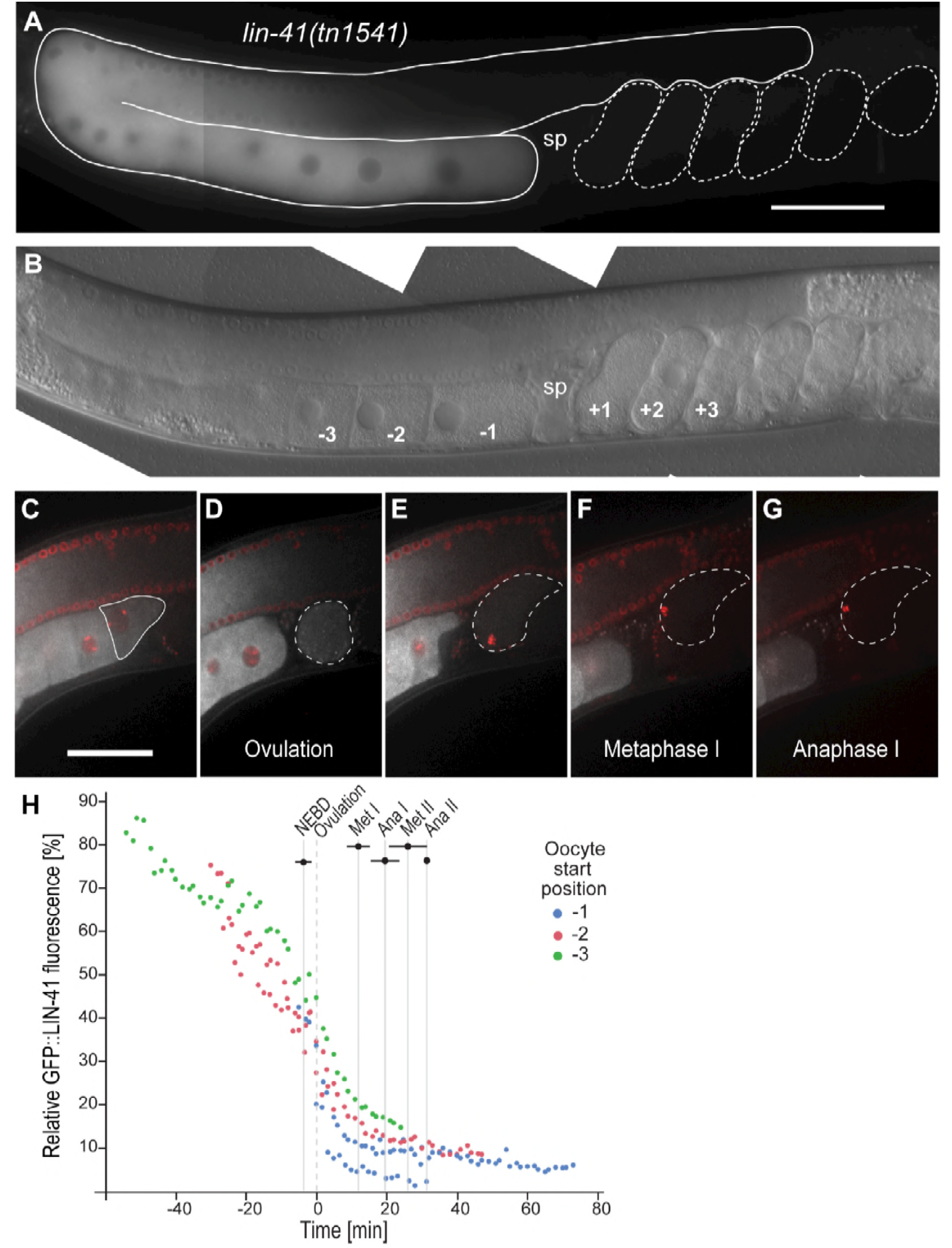
GFP::LIN-41 is eliminated during the first meiotic division. (A, B) Composite GFP (A) and DIC (B) images of a *lin-41(tn1541[gfp::tev::s-tag::lin-41])* adult hermaphrodite. GFP::LIN-41 is apparent in the middle and proximal regions of the germline (solid outline, (A)), with reduced levels in the −1 oocyte immediately adjacent to the spermatheca (sp). The positions of some embryos (dashed outlines, (A)) and oocytes are indicated relative to the spermatheca in (B); a fertilized embryo in the spermatheca would be at the zero position. These labels and naming conventions are used throughout. 100 ms GFP exposures; scale bar, 50 μm. (C–G) Time-lapse images of GFP::LIN-41 (white) and mCHERRY::HISTONE-labeled chromosomes (red) were acquired in a living *lin-41(tn1541); itls37[pie-1p::mCherry:::H2B::pie-l 3’UTR, unc-119(+)]* adult hermaphrodite by confocal microscopy. Images are shown for select time points (*t*) prior to meiotic maturation (C, *t*=-4.5 min), at ovulation (D, *t*=0 min), and during the first meiotic division (E, *t*=+4 min; F, *t*=+11.8 min; G, *t*=+16.9 min) as an individual oocyte (C, solid outline) progresses from the −1 to the +1 position and through the OET (D–G, dashed outlines). Scale bar, 50 μm. Movie SI, worm #1, shows the complete time-lapse series from which the still images were taken. (H) Five oocytes were imaged as they progressed from the −1 position through meiotic divisions; the relative amount of background-corrected GFP::LIN-41 with respect to distal oocytes is shown on the graph at each time point. Three of the oocytes were also imaged at earlier stages as they moved from a more distal location (−2 oocyte (red) or −3 oocyte (green) position) into the −1 oocyte position (blue), as indicated. Timing on the x-axis is relative to ovulation (*t*=0). Bars indicate the standard deviation for different meiotic events (e.g.: NEBD, nuclear envelope breakdown; Met, metaphase; Ana, anaphase).

Here we examine the mechanism by which LIN-41 is eliminated by the end of the first meiotic division. We identify two LIN-41 degradation domains, Deg-A and Deg-B, and a potential CDK-1 phosphorylation site within Deg-A that are individually required for efficient degradation. Transplantation of both LIN-41 degradation domains into OMA-2 results in the premature degradation of the resulting fusion protein during meiosis. Furthermore, we find that a Skp, Cullin, F-box (SCF) E3 ubiquitin ligase complex containing the substrate recognition subunit SEL-10 promotes the degradation of LIN-41 and likely functions through the newly identified degradation domains of LIN-41. SEL-10 is a highly conserved F-box protein important forcell-cycle regulation in both yeast (cell division control protein 4 (Cdc4)) and humans (F-box and WD repeat domain protein (FBW7)) (reviewed in Deshaies and Ferrell 2001; Welcker and Clurman 2008).

Intriguingly, we show that SEL-10 is also important for the degradation of the tumor suppressor protein GLD-1/STAR RNA-binding protein, which is required for oocyte differentiation and represses translation in oocytes (Francis *et al.* 1995a,b; Jones and Schedl 1995; Lee and Schedl 2001; Shumacher *et al. 2005;* Wright *et al.* 2011; Jungkamp *et al.* 2011; Scheckel *et al.* 2012; Farley and Ryder 2012; Doh *et al.* 2013). GLD-1 was independently identified as a target of SEL-10-mediated degradation by Kisielnicka *et al.* (2018) along with CPB-3, a cytoplasmic polyadenylation element (CPE)-binding (CPEB) protein, which is also important for oocyte development (Boag *et al.* 2005; Hasegawa *et al.* 2006). GLD-1 and CPB-3 are degraded during meiotic prophase, as immature oocytes transition from pachytene to diplotene (Kisielnicka *et al.* 2018), considerably earlier than the degradation of LIN-41 during the OET. This is likely due to differences in signaling-mediated regulation; while the degradation of LIN-41 is regulated by activated CDK-1 (Spike *et al.* 2014a and this work), the degradation of GLD-1 and CPB-3 is regulated by the MAP kinase MPK-1 (Kisielnicka *et al.* 2018 and this work). Surprisingly, the ectopic expression of LIN-41 and GLD-1 in *sel-10* mutants has only minor effects on fertility and the expression of mRNAs that are translationally repressed by either LIN-41 or GLD-1 during oogenesis. We suggest that the LIN-41 that persists in the embryos of *sel-10* and certain *lin-41* mutants is likely inactivated by additional post-transcriptional mechanisms that remain to be identified.

## MATERIALS AND METHODS

### Strains

The genotypes of strains used in this study are reported in Table S1. The following mutations were used: LGI - *mex-3(tn1753[gfp::3xflag::mex-3]), air-2(or207ts), unc-13(el091), rŗf-1(pk1417), gld-1(q485), lin-41(tn1487ts), lin-41(tn1541[gfp::tev::s-tag::lin-41], lin-41(tn1541tn1548), Hn-41(tn1541tn1562), lin-41(tn1541tn1571), lin-41(tn1541tn1618), lin-41(tn1541tn1620), slin-41(tn1541tn1622), lin-41(tn1541tn1628), lin-41(tn1541tn1630), lin-41(tn1541tn1635), lin-41(tn1541tn1638), lin-41(tn1541tn1641), lin-41(tn1541tn1643), lin-41(tn1541tn1645), lin-41(tn1541tn1661), lin-41(tn1541tn1663), lin-41(tn1541tn1665), lin-41(tn1541tn1668), lin-41(tn1541tn1684), lin-41(tn1541tn1775), lin-41(tn1767),fog-3(q470),* and *lin-ll(n566).* LGIII - *mpk-l(galllts), emb-30(tn377ts), cdk-l(ne2257ts), orc-1(tn1732[mng::3xflag::orc-1])* and *cul-2(or209ts).* LGIV-*pgl-l(sam37[pgl-lR765S::mTagRFPT::3xflag)* (kindly provided by Dustin Updike), *cks-l(ne549ts),* and *oma-I(zu405te33).* LGV-*spn-4(tn1699[spn-4::gfp::3xflag]), oma-2(te51), oma-2(cp145[mng::3xflag::oma-2]), oma-2(tn1760[mng::3xflag::degA::oma-2]), oma-2(tn1764[mng::3xflag::degA::degB::oma-2]), lon-3(e2175), sel-10(ar41), sel-10(ok1632), sel-10(n1077), him-5(e1490),* and *fog-2(oz40).* LGX–*meg-1(tn1724[gfp::3xflag::meg-1]).* The following rearrangements were used: *hT2[bli-4(e937) Iet-?(q782) qls48]* (I; III) and *nTl[qls51]* (IV; V). The following transgene insertions were used: *axis1498[pie-1p::gfp::gld-1::gld-1 3’UTR, unc-119(+)](*Merritt *et al.* 2008), *itls37[pie-1p::mCherry::H2B::pie-l 3’UTR, unc-119(+)]* (McNally *et al. 2006*), *ozls5[gld-1::gfp]* I (kindly provided by Tim Schedl), *ozis2[gld-1::gfp]* II (Schumacher *et al.* 2005), and *pwis116[rme-2p::rme-2::gfp::rme-2 3’UTR, unc-119(+)]* (Balklava *et al.* 2007).

### Genome editing

Plasmids capable of expressing guide RNAs (gRNAs) that target the *lin-41* gene were generated as described by Arribere *et al.* (2014) from the vector pRB1017 and sequence-specific olignonucleotides. We estimated the efficiency with which each *lin-41* gRNA was able to target the *lin-41(tn1541)* locus by: (1) co-injecting a mixture of the gRNA plasmid (25 ng/μl), the pDD162 plasmid (Dickinson *et al.* 2013), which supplies the Cas9 enzyme (50 ng/μl), and a coinjection marker *(myo-2p::Tdtomato,* 4 ng/μl) into *lin-41(tn1541)* hermaphrodites, (2) culturing individual F1 progeny that expressed the co-injection marker (typically ≤10 F1s from each injected parent), and (3) determining the number of F1s that segregated F2 progeny with a Dpy *lin-41* loss-of-function (If) phenotype. File S1 reports the sequences and estimated efficiencies of the gRNAs we used to generate the *lin-41* deletions and point mutations described in this work; most were relatively effective at targeting *lin-41.*

During the efficiency experiments for *lin-41* gRNAs #10 and #11, we identified *lin-41(tn1541tn1562)* and *lin-41(tn1541tn1571),* respectively, as GFP::LIN-41-positive *lin-41(lf)* mutants that appeared to have relatively large deletions by PCR. All of the other deletions were generated in a targeted manner by co-injecting two or more *lin-41* gRNA plasmids (25 ng/μl each), a single-stranded oligonucleotide (ssODN) repair template (500 nM), the pDD162 plasmid (50 ng/μl), and a co-injection marker *(myo-2p::Tdtomato,* 4 ng/μl) into *lin-41(tn1541)* hermaphrodites. We used gRNAs on each side of the desired deletion that, in most cases, would not produce substrates for Cas9 digestion after the deletion event. Otherwise, silent mutations were included in the repair template to prevent re-cutting. To identify *lin-41(tn1541)* deletion mutants, we individually placed F1 worms expressing the co-injection marker on plates, allowed them to lay eggs, and then used PCR to screen pools of up to 6 F1 worms. Pools that appeared to be strongly positive for the desired deletion band were rescreened by PCR to identify F1 animals that had segregated candidate deletion mutants among their F2 progeny. Mutants were either allowed to become homozygous or were balanced using *hT2[bli-4(e937) Iet-?(q782) qls48]* (I; III). Essentially the same method was used to generate amino acid substitutions in *lin-41.* However, in those experiments we used only one gRNA and repair events were identified using silent mutations that created restriction enzyme recognition sites in each repair template. Screening therefore consisted of PCR followed by a restriction enzyme digestion, and we only pooled 2 F1s in the initial round of screening so that the repair events would be easy to detect. All edited loci were validated by sequencing, and we were able to obtain multiple independent alleles for most targeted deletions and amino acid substitutions. Where possible, two alleles identical to the repair template (but derived from independently injected parents) were saved and assigned allele names. Other, typically imperfect, gene edits were also kept and given allele names if they were informative or potentially useful. Additional information about all of these alleles, as well as detailed genome editing information, including gRNA, repair template, and PCR primer sequences, is provided in File S1.

*oma-2 (tn1760)* and *oma-2(tn1764)* were created using the method described by Dickinson *et al.* (2015) to create *oma-2(cp145).* Indeed, we were careful to replicate *oma-2(cp145)* as closely as possible; we used the same gRNA plasmid (pDD223) and designed our repair templates to closely mimic pDD271, the repair template used to create *oma-2(cp145).* However, instead of using PCR to generate the 3’ homology arms of the repair templates, which contain sequences derived from both *oma-2* and *lin-41,* we synthesized these sequences as gBIocks (Integrated DNA Technologies, Skokie, IL). We minimized the size and complexity of each gBlock by removing introns from the *lin-41*-encoding sequences. *oma-2(tn1760)* and *oma-2(tn1764)* were perfect matches to the desired repairs (repair templates pCS557 and pCS561, respectively). Gene edited alleles were out-crossed to the wild type before analyzing fertility and embryonic lethality. Specific genome editing details are provided in File S1.

### Microscopy

Movies of GFP::LIN-41, mNG::Deg-A::Deg-B::OMA-2, PGL-1::RFP, and mCHERRY::H2B during the OET were obtained using a Marianas 200 Microscopy Workstation (Intelligent Imaging Innovations) built on an AxioObserver Z.l stand (Carl Zeiss, Thornwood, NY) and driven by SIideBook 6.0 software (Intelligent Imaging Innovations, Denver, CO). The imaging was performed using a 40x oil Carl Zeiss Plan-Apochromat objective lens (numerical aperture of 1.4) and an Evolve electron-multiplying charge-coupled device camera (Photometries, Tucson, AZ). The quantification of GFP::LIN-41 in proximal oocytes and embryos relative to distal oocytes was performed using ImageJ software. All of the other images were acquired on a Carl Zeiss motorized Axioplan 2 microscope with a 63X Plan-Apochromat (numerical aperture 1.4) objective lens using a AxioCam MRm camera and AxioVision software (Carl Zeiss). Image quantifications were also performed using AxioVision, version 4.8.2.0. The average intensity of SPN-4::GFP fluorescence was measured in a ~12 μM diameter circle in the anterior cytoplasm of 1-cell and 2-cell embryos in order to avoid the bright puncta of SPN-4::GFP in the posterior; these are likely P granules, as SPN-4 is known to associate with these non-membrane-bound organelles in embryos (Ogura *et al.* 2003). The amount of diffusely cytoplasmic SPN-4::GFP appeared to be similar throughout the embryo during these early stages of embryogenesis and in each of the strains we analyzed. Likewise, the average intensity of GFP::MEX-3 and mNG::OMA-2 fluorescence was measured in a ~10 μM diameter circle in the oocyte cytoplasm. Fluorescence was measured in the oocytes that expressed detectable levels of each fusion protein under our imaging conditions (100 ms and 120 ms for GFP::MEX-3 and mNG::OMA-2, respectively) and were large enough to fit a ~10 μM diameter circle in the oocyte cytoplasm. GFP::MEX-3 was detected in 4 or 5 proximal oocytes in all strains, consistent with previous observations (Tsukamoto *et al.* 2017). mNG::OMA-2 was detected in 5 or 6 proximal oocytes in the *sel-10(ar41)* mutants and in 7 or more proximal oocytes in the control strain.

### RNA interference

Gene-specific RNA interference (RNAi) was performed by feeding *C. elegans* with double-stranded RNA (dsRNA)-expressing *E. coli* (Timmons and Fire 1998) at 22° using the RNAi culture media described by Govindan *et al.* (2006). RNAi clones were obtained from Source BioScience (Nottingham, UK), and the identity of each RNAi clone verified by DNA sequencing. The RNAi clone used for *cul-2(RNAi)* targets the *cul-2* 3’UTR, which may make it less effective at triggering an RNAi response. Exposure to dsRNA-expressing *E. coli* was initiated during the fourth larval stage and GFP::LIN-41 was examined after 1 and 2 days. *cdk-l(RNAi), skr-l(RNAi)* and *sel-10(RNAi)* at least partially prevented the elimination of GFP::LIN-41 after 1 day, with stronger and more penetrant phenotypic effects on Day 2, while *cul-l(RNAi)* only prevented the elimination of GFP::LIN-41 after 2 days of RNAi treatment. All images of RNAi-treated animals were collected on Day 2.

### Western blots

Proteins were separated using NuPage 4-12% Bis-Tris gels or 3-8%Tris-Acetate gels (Invitrogen, Carlsbad, CA) and visualized after western blotting. Blots were blocked with 5% nonfat dried milk. Primary antibodies used to detect proteins were affinity-purified rabbit anti-LIN-41 R214 (1:20,000 dilution) (Spike *et al.* 2014a), guinea pig anti-LIN-41 GP49E (1:4,000 dilution) (Spike *et al.* 2014a), and rabbit anti-GLD-1 (1:3,000 dilution; kindly provided by Sarah Crittenden and Judith Kimble) (Jan *et al.* 1999). Secondary antibodies used for western blots were peroxidase-conjugated donkey anti-guinea pig (1:40,000 dilution) (Jackson ImmunoResearch, West Grove, PA) and anti-rabbit (1:5,000 dilution) (Thermo Scientific, Waltham, MA) antibodies. Detection was performed using SuperSignal West Femto Maximum Sensitivity Substrate (Thermo Scientific).

### Antibody staining

Dissected gonads stained with either the rabbit anti-phospho-histone H3 (SerlO) antibody (1:400 dilution; Millipore, Burlington, MA) or the rabbit anti-RME-2 antibody (1:50 dilution, kindly provided by Barth Grant) (Grant and Hirsh 1999) were fixed in 3% paraformaldehyde for 1 hour, as described (Rose *et al.* 1997). Dissected gonads stained with the rabbit anti-GLD-1 primary antibody (1:150 dilution; Jan *et al.* 1999) were fixed in 1% paraformaldehyde for 10 minutes. Primary antibodies were detected using either Cy3-conjugated goat anti-rabbit or Alexa 488-conjugated donkey anti-rabbit secondary antibodies (1:500 dilutions; Jackson ImmunoResearch).

### Data availability

All strains and newly-created alleles (see Table S1 and File S1) are available upon request. The sequences of gRNAs, repair templates, PCR primers, *lin-41* alleles, and *oma-2* alleles are presented in File S1. Plasmids producing gRNAs and those containing repair templates for genome editing are available upon request. All Sanger sequencing files are available upon request. Supplemental materials are available at Figshare: https://doi.org/.

## RESULTS

### GFP::LIN-41 is eliminated during the first meiotic division

Germline-expressed LIN-41 is restricted to oogenesis and required to prevent premature M-phase entry and to promote growth of developing oocytes (Spike *et al.* 2014a). In the oogenic germlines of adult hermaphrodites, LIN-41 is expressed from mid-pachytene through subsequent stages of oocyte development, with a notable reduction in LIN-41 levels as oocytes initiate meiotic maturation at the end of oogenesis. Essentially the same pattern is seen in the oocytes of *lin-41(tn1541[gfp::tev::s-tag::lin-41])* adult hermaphrodites; these animals carry a *gfp*-tagged allele of *lin-41* and express only GFP-tagged LIN-41 (GFP::LIN-41), yet have essentially wild-type oocyte development and fertility (Figure 1, A and B; Spike *et al.* 2014a). GFP::LIN-41 is always visible in the oocyte immediately adjacent to the spermatheca (−1 oocyte), but is not detectable in most embryos, suggesting that GFP::LIN-41 is eliminated soon after meiotic maturation and ovulation (Figure 1, A and B; Spike *et al.* 2014a). To more precisely determine the stage at which GFP::LIN-41 is eliminated during the OET, we used time-lapse imaging to examine GFP::LIN-41 as oocytes proceed through meiotic maturation, are ovulated into and fertilized in the spermatheca, and complete their meiotic divisions (Movie S1 and Figure 1, C-H). These images show that GFP::LIN-41 levels drop dramatically after meiotic maturation and ovulation (Figure 1, D and H) and that GFP::LIN-41 is essentially undetectable well before the end of the first meiotic division (Figure 1, F and G). We were able to image several oocytes as they moved into the −1 oocyte position from a slightly earlier developmental stage (−2 or −3 oocyte start position) and completed meiosis. Quantification of GFP::LIN-41 levels in these oocytes revealed that GFP::LIN-41 levels also decline during the late stages of oogenesis, albeit in a somewhat less-dramatic fashion than after meiotic maturation (Figure 1H). During meiotic maturation, the oocyte undergoes a cortical cytoskeletal rearrangement prior to ovulation (McCarter *et al.* 1999). During cortical rearrangement, we observed that GFP::LIN-41 began to localize to punctate structures in the oocyte cytoplasm coincident with the onset of its dramatic disappearance (Movie S1). The nature of these punctate structures is unclear; however, they do not appear to be P granules as most of them do not exhibit colocalization with PGL-1::RFP (Figure S1, A-C).

### Deg-A and Deg-B are required to eliminate GFP::LIN-41 from embryos

To identify the amino acid sequences of LIN-41 required for its elimination from early embryos, we generated a series of deletions in the coding region of the GFP::LIN-41-expressing *lin-41(tn1541)* gene using CRISPR-Cas9-based genome editing approaches (see Materials and Methods and File S1 for details). Collectively, these deletions are predicted to remove 95% of the LIN-41 protein and disrupt all known structural domains of LIN-41 (Figure 2, A and B, and File S1). For each mutant, GFP::LIN-41[Δ] expression was examined to determine whether the deleted portion of LIN-41 is necessary for the elimination of GFP::LIN-41 from embryos (Figure 2, D-F, Figure S2, and Figure S3). Null mutations in *lin-41* are sterile, with small, abnormal oocytes, and some hypomorphic alleles of *lin-41* affect the production of high-quality oocytes (Slack *et al. 2000;* Spike *et al.* 2014a). Thus, GFP::LIN-41[Δ] was often examined in both heterozygotes *(Hn-41(tn1541Δ)/unc-13(el091) Iin-ll(n566)* genotypes) and homozygotes *(lin-41(tn1541Δ)* genotypes), particularly when the *lin-41(tn1541Δ)* homozygotes produced obviously small or abnormal oocytes or produced a significant number of dead embryos (Figure S2, Figure S3, and Table 1). This approach enabled us to determine that two non-overlapping regions in the N-terminal third of LIN-41 are required for the elimination of LIN-41 from embryos (Figure 2B). We will refer to these regions as the LIN-41 degradation domains Deg-A and Deg-B.

**FIGURE 2.**
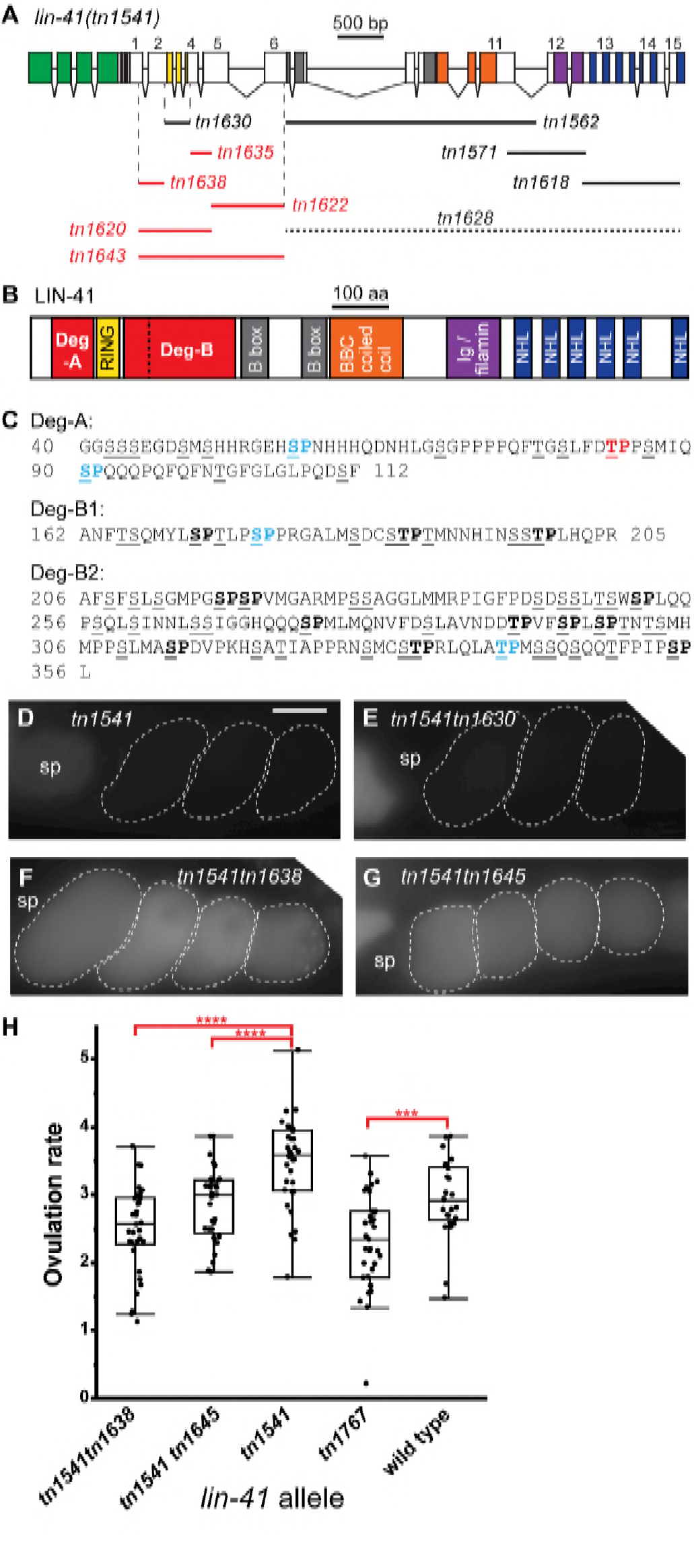
GFP::LIN-41 elimination requires two non-overlapping regions of LIN-41 and a potential phosphorylation site. (A) The exon-intron structure and deletion analysis of *lin-41(tn1541).* Colored boxes indicate exonic regions that encode GFP (green) or previously described protein domains of LIN-41 (see (B)). Deletions made in the context of *lin-41(tn1541)* are drawn as lines, labeled with a deletion-specific allele name, below LIN-41-encoding exons and introns (exons labeled 1-15). GFP::LIN-41 can be detected in the germline of most deletion mutants (solid lines), with one exception *(tn1628,* dotted line). Deletions in red prevent the elimination of GFP::LIN-41 from early embryos. The vertical dashed lines delimit the beginning of Deg-A and the end of Deg-B, respectively. (B) The previously described (RING (yellow), B-box (gray), BBC (orange), Ig/filamin (purple), NHL (blue)) and newly-identified (Deg (red)) protein domains of LIN-41. The vertical dashed line in (B) indicates the two parts of Deg-B, B1 and B2, which are individually removed in *lin-41(tn1541tn1635)* and *lin-41(tn1541tn1622),* respectively. (C) The amino acid sequences of Deg-A, Deg-Bl and Deg-B2. Many of the amino acids are serines and threonines (underlined); some are potential targets of proline-directed serine/threonine [S/T] kinases (bold) and have had the [S/T] residue changed to an alanine (colored and bold) in the context of *lin-41(tn1541).* The T83A mutation in Deg-A results in the persistence of GFP::LIN-41[T83A] in embryos (red), whereas the other changes do not (indicated in blue font). (D-G) GFP::LIN-41 is eliminated from the early embryos (dashed outlines) of *lin-41(tn1541)* (D, control) and *lin-41(tn1541tn1630)* (E, RING deleted) homozygous mutants but persists in the early embryos of *lin-41(tn1541tn1638)* (F, Deg-A deleted) and *lin-41(tn1541tn1645)* (G, LIN-41[T83A]) homozygous mutants. The position of the spermatheca (sp) is indicated, for reference. 100 ms GFP exposures; scale bar, 20 μm. (H) The rate of ovulation is slightly reduced in mutants with a compromised LIN-41 Deg-A domain. Ovulation rate is expressed as the number of ovulations/gonad arm/hour and was measured in at least 25 Day 2 adults. Significance was determined using a Student’s *t*test: P<.001 is indicated by 3 asterisks, P<.0001 is indicated by4 asterisks. *itls37[pie-1p::mCherry:::H2B::pie-l 3’UTR, unc-119(+)]* was also present in each of the GFP::LIN-41-e×pressing strains; it is not expected to alter the ovulation rate.

**Table 1.** Fertility and fecundity of *lin-41* alleles at 20°C

| GENOTYPE | PREDICTED | FERTILE | BROOD | DEAD |
| --- | --- | --- | --- | --- |
|  | PROTEIN | (%)<sup>a</sup> | SIZE<sup>b</sup> | EMBRYOS |
|  | CHANGE |  |  | (%)<sup>c</sup> |
| *lin-41(tn1541)* | N-terminal | 100 | 316 ± 39<sup>d</sup> | 0.3 |
|  | GFP | (n=68) | (n=6) | (n=361) |
| *lin-41(tn1541tn1618)<sup>e,f</sup>* | Δ NHL | 1.5 | 1 | ND |
|  | (AA 819-1128) | (n=65) | (n=1) |  |
| *lin-41(tn1541tn1571)<sup>e,f</sup>* | Δ Ig | 78.5 | 11 ± 12 | 57.1<sup>g</sup> |
|  | (AA 677-824) | (n=65) | (n=9) | (n=35) |
| *lin-41(tn1541tn1562)<sup>e,f</sup>* | Δ Bbox-CC | 84 | 6 ± 3 | ND |
|  | (AA 356-707)<sup>h</sup> | (n=87) | (n=17) |  |
| *lin-41(tn1541tn1643)<sup>e</sup>* | Δ N-terminal | 66 | 6 ± 4 | 75.4<sup>g</sup> |
|  | (AA 40-356) | (n=90) | (n=48) | (n=142) |
| *lin-41(tn1541tn1620)<sup>e</sup>* | Δ N-terminal | 97 | 39 ± 32 | 36.4<sup>g</sup> |
|  | (AA 40-205) | (n=67) | (n=10) | (n=110) |
| *lin-41(tn1541tn1622)<sup>e</sup>* | Δ Deg-B2 | 100 | 33 ± 16 | 39.0<sup>g</sup> |
|  | (AA 206-356) | (n=65) | (n=6) | (n=105) |
| *lin-41(tn1541tn1635)* | Δ Deg-B1 | 100 | 127 ± 108 | 2.9 |
|  | (AA 162-205) | (n=70) | (n=10) | (n=105) |
| *lin-41(tn1541tn1630)* | Δ RING | 98.5 | 210 ± 87 | 2.8 |
|  | (AA 113-161) | (n=65) | (n=12) | (n=144) |
| *lin-41(tn1541tn1638)* | $\Delta$ Deg-A | 100 | 217 ± 103 | 6.3 |
|  | (AA 40-112) | (n=70) | (n=10) | (n=174) |
| *lin-41(tn1541tn1645)* | T83A | 100 | 251 ± 86 | 1.0 |
|  |  | (n=70) | (n=10) | (n=193) |
| *lin-41(tn1767)* | T83A | 98.3 | 313 ± 31 | 0.0 |
|  |  | (n=120) | (n=6) | (n=176) |

<sup>a</sup> Fertile animals produced at least 1 viable offspring.
<sup>b</sup> The average number of progeny that hatched from fertile animals ± the standard deviation.
<sup>c</sup> The percent lethality among the embryos laid on Day 1 of adulthood.
<sup>d</sup> Essentially identical to the *lin-41(tn1541)* brood size previously reported in Spike *et al.* 2014 (319 ± 28 (n=30)).
<sup>e</sup> The progeny of *lin-41/hT2[q/s48]* hermaphrodites.
<sup>f</sup> These animals have a dumpy (Dpy) body shape, as previously described for *lin-41(lf)* alleles (Slack *et al.* 2000).
<sup>g</sup> Some of the embryos laid were small or otherwise appeared to be abnormal.
<sup>h</sup> The minimum number of amino acids removed by *tn1562*. Assuming the use of an in-frame 5' splice site in the 17-bp insertion, either one amino acid (L) or five amino acids (LSPLL) would replace amino acids 356-707.

The LIN-41 Deg-A domain is defined by the *lin-41(tn1541tn1638)* deletion allele. This deletion is predicted to affect GFP::LIN-41 by removing 73 amino acids on the N-terminal side of the LIN-41 RING domain (Figure 2C and File S1). *lin-41(tn1541tn1638)* is immediately adjacent to, but does not overlap, the *Hn-41(tn1541tn1630)* deletion, which is predicted to affect GFP::LIN-41 by removing the LIN-41 RING finger domain (see File S1 for deleted residues). Consistent with previous amino acid substitution and transgenic rescue data (Tocchini *et al.* 2014), the RING domain is not required for the elimination of GFP::LIN-41 from embryos (Figure 2E and Figure S2). The LIN-41 Deg-B domain is defined by two contiguous, but non-overlapping, deletions on the C-terminal side of the LIN-41 RING domain. The *lin-41(tn1541tn1635)* deletion is predicted to affect GFP::LIN-41 by removing 44 amino acids on the C-terminal side of the LIN-41 RING domain (Deg-Bl) (Figure 2C and File S1). Compared to the Deg-A deletion mutant *(lin-41(tn1541tn1638)),* the Deg-Bl deletion mutant *Ųin-41(tn1541tn1635))* has a relatively low but detectable level of GFP::LIN-41[Δ] in early embryos (compare Figure S2, K and M). However, the *lin-41(tn1541tn1622)* deletion, which defines Deg-B2, and the remaining 151 amino acids of Deg-B (Figure 2C and File S1), has a robust defect in the elimination of GFP::LIN-41[Δ] from early embryos that is apparent in both heterozygous and homozygous deletion mutants (Figure S2, E and G). Finally, deletions predicted to affect GFP::LIN-41 by removing amino acids and structural domains C-terminal to Deg-B were able to eliminate GFP::LIN-41[Δ] from early embryos (Figure 2A and Figure S3). Interestingly, we found that C-terminal domains could only be removed individually or in small groups, as GFP::LIN-41[Δ] was not detectable when a majority of the C-terminus wasremoved *(lin-41(tn1541tn1628)* deletion; Figure 2A and Figure S3, M-P).

### LIN-41[T83] is required to eliminate LIN-41 from embryos

The results described above indicate that the elimination of GFP::LIN-41 does not depend on any of the previously described structural domains of LIN-41, but instead requires two new regulatory domains. Analysis of the amino acid sequences of Deg-A and Deg-B shows that each regulatory domain contains many possible phosphorylation sites (Figure 2C). Previously published results indicate that the elimination of GFP::LIN-41 from embryos also requires CDK-1 (Spike *et al.* 2014a), a highly conserved proline-directed serine/threonine kinase essential for M-phase entry during oocyte meiotic maturation in *C. elegans* (Boxem *et al.* 1999). Thus, we hypothesized that LIN-41 might be a direct target of CDK-1 activity, and that phosphorylation of either Deg-A or Deg-B by CDK-1 could be sufficient to trigger the elimination of GFP::LIN-41 from embryos. 18 minimal CDK-1 consensus sequences ([S/T]P) are present in Deg-A and Deg-B (Figure 2C), but only a single site, found in Deg-Bl, conforms to an expanded CDK1 consensus sequence ([S/T]PX[K/R]) (Ubersax *et al.* 2003). However, changing the potentially phosphorylated residue at this site to an alanine (S176A; e.g.: *lin-41(tn1541tn1641))* had no effect on the elimination of GFP::LIN-41 from embryos (Figure 2C and Figure S4A).

We therefore shifted our focus to Deg-A, which is relatively small and contains only three potential CDK-1 target sites, but strongly prevents the elimination of GFP::LIN-41 from embryos (Figure S2M). Each site in Deg-A was tested individually to see if it is required for the elimination of GFP::LIN-41 from embryos. Although the mutations S57A and S90A (e.g.: *lin-41(tn1541tn1663)* and *lin-41(tn1541tn1661),* respectively) had no discernable effect (Figure S4, C and E), the T83A mutation (e.g.: *lin-41(tn1541tn1645))* strongly prevented the elimination of GFP::LIN-41 from embryos, similar to the Deg-A deletion mutant (Figure 2G, Figure S2M, and Figure S4G). Time-lapse imaging ofoocyte meiotic maturation, ovulation, and fertilization documents that the T83A mutation strongly abrogates the elimination of GFP::LIN-41 (Movie S2). During cortical rearrangement, GFP::LIN-41[T83A] localized partially to dynamic punctate structures like GFP::LIN-41 (compare Movie S1 and Movie S2); however unlike the wild-type protein, puncta of GFP::LIN-41[T83A] were also observed during the meiotic divisions. Furthermore, GFP::LIN-41[T83A] persisted through multiple embryonic cleavage divisions and became at least partially associated with P granules by the 2-cell stage (Movie S2 and Figure SI, D-l). These results are consistent with the possibility that phosphorylation of LIN-41 by a proline-directed S/T kinase, such as CDK-1, promotes the rapid elimination of GFP::LIN-41 upon the onset of meiotic maturation. As a further assessment, we replaced T83 with a glutamic acid residue (T83E) (e.g.: *lin-41(tn1541tn1684)),* which is negatively charged and might function as a phosphomimetic. However, T83E did not result in the premature elimination of GFP::LIN-41, as when CDK-1 is prematurely activated (Spike *et al.* 2014a). Instead, T83E prevented the elimination of GFP::LIN-41 from embryos, similarto T83A (Figure S4, G and I). Thus, we conclude that either T83E does not effectively mimic phosphorylation at this particular site or T83 might not be phosphorylated. In fact, phosphorylation sites that function to recruit adapter proteins are of ten not recognized by binding partners after phosphomimetic substitution (Dephoure *et al.* 2013), and this is a possible explanation for the function of T83 and the Deg domains of LIN-41, as we will describe.

A requirement for the extreme N-terminus of LIN-41 (amino acids 1-39) with respect to the elimination of GFP::LIN-41 was not examined in the *lin-41(tn1541)* deletion analysis. Genetic analysis suggests that this region of LIN-41 is important for down-regulating *lin-41* function specifically in the male tail (Del Rio-Albrechtsen *et al.* 2006). Gain-of-function (gf) alleles that affect this part of LIN-41 have a defect in male tail tip retraction, while hermaphrodites appear overtly wild-type. *lin-41(tn1541)* males also have a male tail tip retraction defect (Figure S5, A and B), suggesting that the GFP tag on the N-terminus of LIN-41 disrupts this male-specific function. Furthermore, the amino acid change found in the *lin-41(bx37gf)* allele (G35R) does not affect the elimination of GFP::LIN-41 from early embryos *(lin-41(tn1541tn1665);* Figure S5, Cand E). For these reasons, we suspectthatthe extreme N-terminus of LIN-41 is unlikely to be involved in the elimination of GFP::LIN-41 from early embryos. One possibility, however, might be that the N-terminal GFP tag on GFP::LIN-41 compromises a function that is required redundantly with Deg-A or Deg-B. To explore this possibility, we generated worms expressing LIN-41[T83A] *(lin-41(tn1767))* and asked whether the untagged protein also persists in embryos. Using western blots, we found that LIN-41 was undetectable in a lysate made from wild-type embryos, but that LIN-41[T83A] was clearly present in a lysate prepared from *lin-41(tn1767)* mutant embryos (Figure S6C). Thus, the T83A mutation abrogates the elimination of both LIN-41 and GFP::LIN-41 from embryos.

### Functional requirements for individual LIN-41 domains

LIN-41 is a large protein with two well-defined domains that are proposed to have strikingly different activities. The first of these is actually a multi-domain grouping called a <u>TRI</u>partite <u>M</u>otif (TRIM) that contains RING, B-box, and coiled-coil (CC) domains; many TRIM proteins are thought to function as RING finger E3 ubiquitin ligases (Ikeda and Inoue 2012). The second functional domain is an RNA-binding domain composed of 6 NHL (<u>N</u>CL-1, <u>H</u>T2A and <u>L</u>IN-41) repeats at the C-terminus of LIN-41 (Slack and Ruvkun 1998; Loedige *et al.* 2015; Kumari *et al.* 2018). Forward and reverse genetic analyses strongly indicate that the NHL domain is important for both the germline and somatic functions of *C. elegans* LIN-41 (Slack *et al.* 2000; Spike *et al.* 2014a; Tocchini *et al.* 2014), consistent with the identification of LIN-41 as a translational repressor in both tissue types (Spike *et al.* 2014b; Aeschimann *et al.* 2017; Tsukamoto *et al.* 2017). By contrast, a deletion of the entire LIN-41 RING domain (Figure 2A), which confers *in vitro* E3 ligase catalytic activity to mouse LIN41 and other TRIM proteins (Rybak *et al.* 2009; Esposito *et al.* 2017), results in appreciable fertility (brood size of 210 ± 87; Table 1) and thus is non-essential for *C. elegans* oogenesis. As described below, the phenotypes seen in *lin-41(tn1541)* deletion mutants are consistent with prior observations and provide additional insights into the functions of LIN-41 protein domains.

#### TRIM (Ring, B-box, CC) domain

Deletion of the RING finger in the context of GFP::LIN-41 (GFP::LIN-41[ΔRING]) results in only mild defects. Most *lin-41(tn1541tn1630)* animals are fertile and have a large number of progeny; no strong defects in oogenesis, embryonic development or body shape are evident (Table 1 and Figure S2, I and J). We did note, however, that *lin-41(tn1541tn1630)* animals appear to be slightly sick and that they produce ~33% fewer progeny than *lin-41(tn1541)* hermaphrodites (Table 1). Interestingly, deletion of the other two TRIM sub-domains (GFP::LIN-41[ΔBbox-CC]) causes a much stronger reduction in LIN-41 function. Most (84%) *lin-41(tn1541tn1562)* hermaphrodites are fertile, but produce very few progeny (6 ± 4) and have obvious defects in oogenesis as well as a Dumpy (Dpy) body shape (Table 1; Figure S3, A and B). Thus, *lin-41(tn1541tn1562)* is clearly a hypomorphic allele of *lin-41* that affects both its germline and somatic functions.

We note that *lin-41(tn1541tn1562)* might remove additional residues beyond the Bbox-CC region because, unlike the other *lin-41(tn1541Δ)* mutants we created, *lin-41(tn1541tn1562)* is not a precise exon-exon fusion and requires a new in-frame splicing event to make a full-length protein (Figure 2A and File S1). However, the deletion in this mutant was accompanied by the insertion of a small sequence that includes two potential 5’ splice site consensus sequences; both are in-frame with the downstream exon. Furthermore, the relative size of GFP::LIN-41[ΔBbox-BBC] on SDS-PAGE western blots is consistent with what we expect to see for the protein made by this particular deletion mutant (File S1 and Figure S6B). This is also true for the other GFP::LIN-41[Δ] proteins we detected on western blots using anti-LIN-41 antibodies (Figure S6, A and B).

#### IG/filamin domain

IG/filamin (IG) domains are only found in a subset of TRIM-NHL proteins; structural analysis of this part of *C. elegans* LIN-41 suggests that it forms a classic IG-Iike domain fold (Tocchini *et al.* 2014). The IG domain has been proposed to function, along with the coiled-coil domain, as a binding platform for proteins that repress the translation of NHL-bound target mRNAs (Loedige *et al.* 2013). *lin-41(ma104)* is a hypomorphic allele that likely disrupts the structure and function of the LIN-41 IG domain (Tocchini *et al.* 2014). As previously reported (Spike *et al.* 2014b), outcrossed *lin-41(ma104)* mutant hermaphrodites have mild oocyte defects and a reduced, but still substantial, brood size of 181 progeny (n=12). Deletion of the IG domain in the context of GFP::LIN-41 (GFP::LIN-41[ΔIG]) results in stronger defects. Most *lin-41(tn1541tn1571)* hermaphrodites are fertile, with a very low brood size (11 progeny) and obvious defects in oogenesis (Table 1 and Figure S3, C and D). Both alleles also result in worms with an obviously Dpy body shape. Thus, despite the difference in brood size, the alleles that affect the IG domain are hypomorphic and reduce both the germline and somatic functions of *lin-41.* Indeed, it is potentially misleading to conclude that the relative severities of *lin-41(ma104)* and *lin-41(tn1541tn1571)* are meaningful, as LIN-41 function may be slightly compromised in the *lin-41(tn1541)* mutant despite its wild-type brood size (316 ± 39; Table 1; Spike *et al.* 2014a). For example, the introduction of the LIN-41[D1125N] amino acid change that results in a temperature-sensitive (ts) phenotype in an otherwise wild-type LIN-41 protein (e.g.: *lin-41(tn1487(ts);* 100% fertile at 15°C (n=224), average brood size of 104 (n=64); Spike *et al.* 2014a) results in a stronger, but still hypomorphic, phenotype in a GFP::LIN-41 mutant background (e.g.: *lin-41(tn1541tn1548))* 71% fertile at 15°C (n=21), average brood size of 3 (n=15)).

#### The NHL domain

Deletion of the C-terminal NHL domain in the context of GFP::LIN-41 (GFP::LIN-41[ΔNHL]) results in a strong loss-of-function *lin-41* phenotype. Whereas *lin-41(tn1541)* hermaphrodites are fertile, with normal oocyte development and overall appearance, nearly all (98.5%) *Hn-41(tn1541tn1618)* hermaphrodites are sterile and have a Dpy body shape (Table 1). Oogenesis is extremely abnormal in most animals (Figure S3, G and H), although *lin-41(tn1541tn1618)* hermaphrodites appear capable of producing embryos on occasion (Table 1 and Figure S3, I and J). The fact that deletion of the LIN-41 NHL domain does not result in 100% sterility is surprising because the *lin-41(n2914)* null mutation has never been observed to produce progeny. Thus, LIN-41 can exhibit some, albeit very low, biological function in the absence of the NHL domain. We suggest, that this low-level function may be mediated through components of the LIN-41 RNP (Spike *et al.* 2014b; Tsukamoto *et al.* 2017). We confirmed that CDK-1 exhibits premature activation in post-dauer *lin-41(tn1541tn1618)* mutants, as it does in *lin-41(n2914)* null mutants, by staining adult hermaphrodite germlines with an antibody specific to histone H3 phosphorylated on Serine 10 (pH3(S10)). This antibody stains the nucleoplasm and condensed chromosomes of wild-type diakinesis-stage oocytes as they prepare enter M-phase nearthe spermatheca (Hsu *et al.* 2000). Both M-phase and anti-pH3(S10) staining occur prematurely in strong loss-of-function *lin-41* mutants and are *cdk-1-* dependent (Spike *et al.* 2014a). As expected for a strong loss-of-function mutant, and consistent with the idea that premature M-phase entry and CDK-1 activation occur prematurely, we detected pH3(S10)-positive condensed chromosomes in or near the loop region of the gonad, and just after the end of pachytene, in most *lin-41(tn1541tn1618)* oogenic germlines (n=6/9). Interestingly, GFP::LIN-41[ΔNHL] forms abnormal aggregates in the oocytes of *Hn-41(tn1541tn1618)* homozygotes; these aggregates are not seen in heterozygotes (Figure S3, G, I, and K). The reason for this aberrant pattern of localization is unknown, but GFP::LIN-41[ΔNHL] aggregation is also seen in post-dauer *lin-41(tn1541tn1618); cdk-l(RNAi)* animals (n=32), and therefore does not depend on the dysregulation of *cdk-1* function that occurs during oogenesis in strong loss-of-function *lin-41* mutants (Spike *et al.* 2014a). These aggregates may reflect abnormal biogenesis of LIN-41 RNPs in the absence of the NHL domain.

#### Meiotic degradation domains are nonessential

We initially hypothesized that the deletion of LIN-41 degradation domains might result in a gain-of-function phenotype that would impact fertility or embryonic viability. However, *lin-41(tn1541tn1643),* a large deletion that removes Deg-A, the RING finger, and Deg-B in the context of GFP::LIN-41, behaves as a recessive hypomorph that preferentially affects germline function. Homozygous mutants do not have a strong Dpy phenotype, but do have an extremely small brood size and display obvious defects in oogenesis and embryogenesis (Table 1 and Figure S2, 0 and P). In contrast, heterozygous *lin-41(tn1541tn1643)* mutants appear essentially normal (n=20). This is also true for deletions that subdivide the large N-terminal region of LIN-41, such as *lin-41(tn1541tn1620)* and *lin-41(tn1541tn1622)* (Table 1 and Figure S2, A-H). Indeed, even when homozygous, the relatively small Deg-A deletion *(lin-41(tn1541tn1638)),* which results in abundant GFP::LIN-41[ΔDeg-A] in early embryos, appears to have minimal consequences for GFP::LIN-41 function at 20° (Table 1 and Figure S2, M and N). Likewise, animals expressing LIN-41[T83] and GFP::LIN-41[T83] appear essentially wild-type; the latter have only a slightly reduced brood size relative to GFP::LIN-41-expressing controls (Table 1 and Figure S4, G and H). Consequently, we decided to look carefully at the ovulation rates of the minimally affected LIN-41 Deg-A deletion *(lin-41(tn1541tn1638))* and T83A point mutants *(lin-41(tn1541tn1645)* and *lin-41(tn1767)).* Oocyte meiotic maturation is a rate-limiting step for hermaphrodite fertility and the ovulation rate approximates the rate ofoocyte meiotic maturation (McCarter *et al.* 1999; Miller *et al.* 2001, Govindan *et al.* 2006). Importantly, several aspects of nuclear and cytoplasmic oocyte maturation occur prematurely in *lin-41(lf)* mutations (Spike *et al.* 2014a,b;Tsukamoto *et al.* 2017). Deg-Adomain mutants exhibit mean ovulation rates that are significantly reduced relative to genotype-matched controls (Figure 2H). Interestingly, the mean ovulation rate of the *lin-41(tn1541)* control strain was elevated relative to wild-type animals (Figure 2H, 3.4 vs 2.9 ovulations/gonad arm/hr). Together, these observations suggest that (1) *lin-41(tn1541)* might be a weak hypomorph that causes a slight increase in the oocyte maturation rate and (2) Deg-A domain mutants cause the opposite phenotype, a reduced oocyte maturation rate. These changes in the rate of oocyte maturation are relatively modest, however, and our phenotypic analyses generally suggest that the elimination of LIN-41 from early embryos is not a critical control point for regulating LIN-41 function or activity levels *in vivo.*

### LIN-41[Deg] domains are sufficient for degradation

OMA-1 and OMA-2 (OMA-1/2) are functionally redundant cytoplasmic RNA binding proteins expressed in oocytes and early embryos (Detwiler *et al.* 2001) that co-purify with LIN-41 RNP complexes (Spike *et al.* 2014b, Tsukamoto *et al.* 2017). OMA-1/2 levels remain high in 1-cell embryos until the first mitotic division, when they are rapidly degraded (Lin 2003; Nishi and Lin 2005; Shirayama *et al.* 2006; Stitzel *et al.* 2006). The expression and subsequent elimination of OMA-2 can be easily visualized in *oma-2(cp145)* mutants (Dickinson *et al.* 2015; Figure 3A and Figure S7, A and B), which express an mNeonGreen-tagged form of OMA-2 that is largely functional *in vivo* (Table 2, compare *oma-l(zu405te33); oma-2(cp145)* to *oma-l(zu405te33); oma-2(te51)* and *oma-l(zu405te33)).* Based on these attributes, we decided to test whether the LIN-41 Deg domains are sufficient to induce the premature degradation of mNG::OMA-2 during meiosis. Molecularly, we chose to place LIN-41 DEG domains between mNeonGreen and OMA-2 (Figure 3, A-C), as this is similar to their locations in GFP::LIN-41 (Figure 2, A and B) and no structural (e.g., X-ray crystallographic) data are available to aid the experimental design. Using the same method that Dickinson *et al.* (2015) used to make *oma-2(cp145),* we created two new *oma-2* alleles that also contain *lin-41*-encoded Deg domains and examined the pattern of mNG::DEG::OMA-2 accumulation prior to the first mitotic division (Figure 3 and Figure S7).

**FIGURE 3.**
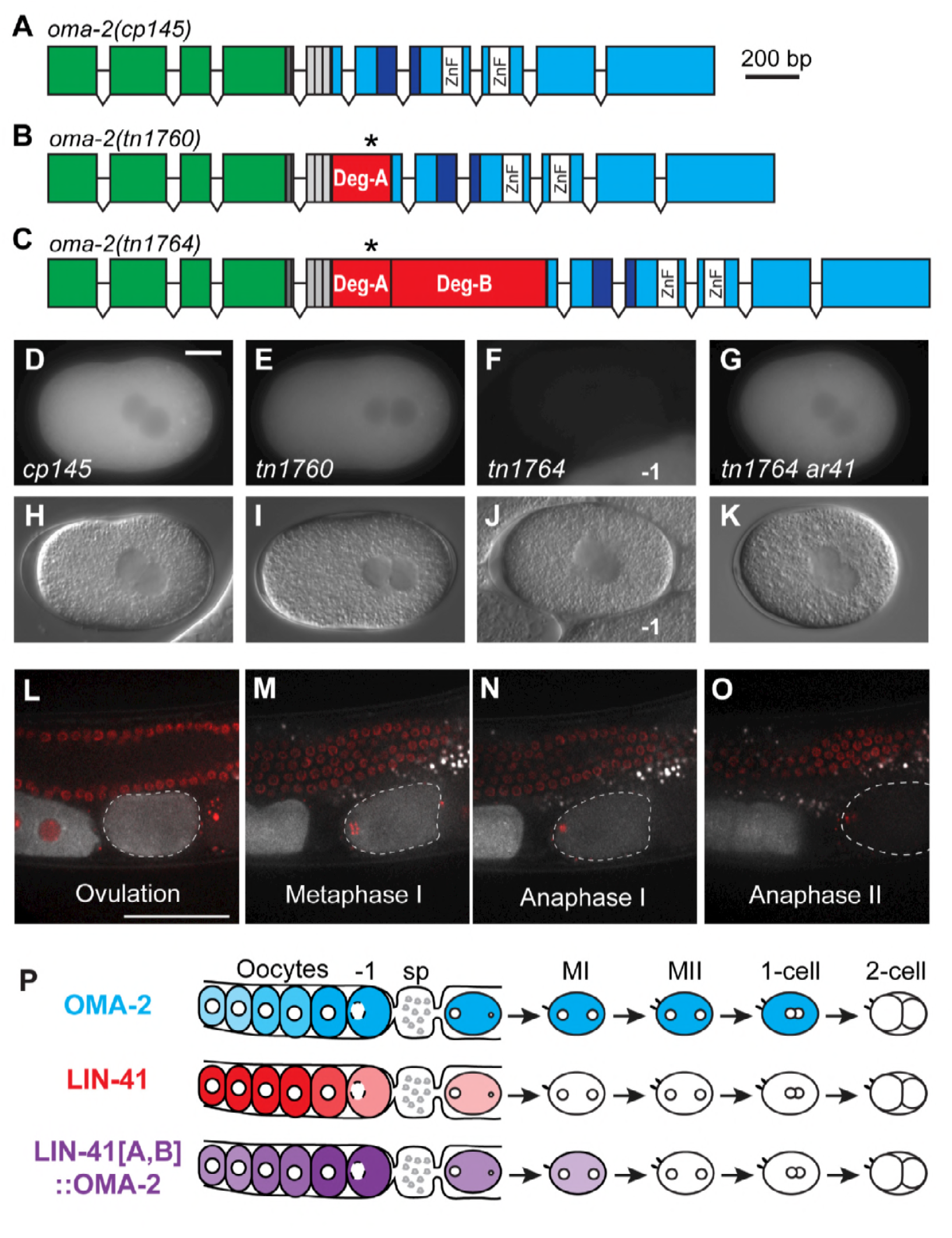
LIN-41 degradation domains when implanted into mNG::0MA-2 promote its rapid elimination during meiosis. (A–C) The exon-intron structures of *oma-2(cp145[mng::tev::3xflag::oma-2]), oma-2(tn1760[mng::tev::3xflag::deg-a::oma-2])* and *oma-2(tn1764[mng::tev::3xflag::deg-a::deg-b::oma-2]).* Boxes represent exonic regions that encode mNeonGreen (green), the tobacco etch virus cleavage site (TEV, dark gray), FLAG epitope tags (light gray), LIN-41 Deg-A and Deg-B domains (red), the likely TAF-4 interaction domain of OMA-2 (dark blue), two OMA-2 CCCH zinc fingers (white), and other OMA-2 coding sequences (cyan). The position of LIN-41 T83 within the LIN-41 Deg-A domain is indicated by an asterisk. (D–K) GFP (D–G) and DIC (H–K) images of *oma-2(cp145)* (D,H), *oma-2(tn1760)* (E,l), *oma-2(tn1764)* (F, J) and *oma-2(tn1764) lon-3(e2175) sel-10(ar41)* (G, K) 1-cell embryos at pronuclear meeting (E, I), orjust slightly later, as the pronuclei begin a counter-clockwise rotation (D, F–G, H, J, and K) prior to NEBD and the first mitotic division. Part of a −1 oocyte is visible in (F, J) and is indicated for reference. 150 ms GFP exposures; scale bar, 10 μm. (L-O) Time-lapse images of mNG::Deg-A,B::OMA-2 (white) and mCHERRY::HISTONE-labeled chromosomes (red) were acquired in a living *oma-2(tn1764); itls37[pie-1p::mCherry:::H2B::pie-l 3’UTR, unc-119(+)]* adult hermaphrodite by confocal microscopy. Images are shown for select time points (*t*) at ovulation (L, *t*=0 min), during the first (M, *t*=+5 min, N, *t*=+10.5 min) and second meiotic divisions (0, *t*=+24.5 min) as an embryo (dashed outline) progresses through both meiotic divisions. See Movie S3 for the complete time-lapse sequence. Scale bar, 50 μm. (P) A visual summary of the dynamic expression patterns of mNG::OMA-2 (cyan), GFP::LIN-41 (red) and mNG::Deg-A,B::OMA-2 (purple). Oocytes are to the left and embryos are to the right of the spermatheca (sp). Meiotic embryos (Ml, Mil) have completed their respective divisions.

**Table 2.** Sterility and embryonic lethality in *oma-2* and *oma-1; oma-2* mutant strains at 20°C

| Genotype | Embryos laid<sup>a</sup> | Dead embryos (%) |
| --- | --- | --- |
| *oma-2(cp145)* | 314 ± 48 (n=6) | 0.6 (n=1793) |
| *oma-2(tn1760)* | 306 ± 39 (n=6) | 0.6 (n=1835) |
| *oma-2(tn1764)* | 300 ± 45 (n=6) | 2.1 (n=1797) |
| *oma-2(tn1764) lon-3(e2175) sel-10(ar41)* | 288 ± 26 (n=6) | 1.1 (n=1727) |
| *oma-1(zu405te33)* | 261 ± 18 (n=6) | 0.8 (n=1568) |
| *oma-1(zu405te33); oma-2(te51) M+Z<sup>-b</sup>* | 0 (n>21)<sup>c</sup> | NA |
| *oma-1(zu405te33); oma-2(cp145)* | 212 ± 29 (n=12) | 12.3<sup>d</sup> (n=2516) |
| *oma-1(zu405te33); oma-2(tn1760)* | 224 ± 35 (n=6) | 60.3 (n=1341) |
| *oma-1(zu405te33); oma-2(tn1764) M+Z<sup>-b,e</sup>* | 249 ± 29 (n=5) | 100 (n>1256) |
| *oma-1(zu405te33); oma-2(tn1764) lon-3(e2175) sel-10(ar41) M+Z<sup>-b,f</sup>* | 246 ± 32 (n=6) | 89.4 (n=1476) |
| *oma-1(zu405te33); oma-2(tn1764) lon-3(e2175) sel-10(ar41)* | 196 ± 56 (n=6) | 84.9 (n=1175) |

<sup>a</sup> Average number of embryos laid per worm ± standard deviation.
<sup>b</sup> M+Z<sup>-</sup> animals were the progeny of *nT1[qIs51]* balancer-containing parents, which are heterozygous for both *oma-1* and *oma-2*. All other animals were the progeny of parents of the listed genotype.
<sup>c</sup> Sterile, with a defect in meiotic maturation as described by Detwiler *et al.* 2001.
<sup>d</sup> Percent embryo lethality was variable among the 12 parents analyzed; it ranged between 6 and 35%.
<sup>e</sup> These animals lay many eggs, none of which hatch (n=30).
<sup>f</sup> These animals lay many eggs, some of which hatch (n=24).

We began by testing LIN-41 Deg-A, which contains the possible CDK-1 target site (T83) required for the elimination of LIN-41. *oma-2(tn1760)* mutants express mNG::Deg-A::OMA-2 in oocytes and 1-cell embryos. Similar to mNG::OMA-2, this protein is present in 1-cell embryos just prior to the first mitotic division but eliminated from older embryos (Figure 3, D and E, and Figure S7, A, B, E, and F). Indeed, the only obvious difference was the amount of mNG in older 1-cell pronuclear stage embryos, which was significantly reduced in *oma-2(tn1760)* embryos compared to *oma-2(cp145)* controls (compare Figure 3, D and E, and Figure S8A). This reduction might be caused by Deg-A-mediated destabilization of the mNG::OMA-2 fusion protein (see below), but is not equivalent to the rapid elimination of GFP::LIN-41 that occurs in meiosis I (Movie S1 and Figure 1). Because Deg-A on its own was not sufficient to trigger the rapid elimination of mNG::OMA-2 in 1-cell embryos, we tested LIN-41 Deg-A and Deg-B together. *oma-2(tn1764)* mutants express mNG::Deg-A, Deg-B::OMA-2 in oocytes, but in 1-cell embryos the amount of mNG is substantially reduced or absent (Figure 3F and Figure S7, G and I). To more precisely determine the stage at which mNG::DEG-A, Deg-B::OMA-2 is eliminated, we used time-lapse imaging (Movie S3 and Figure 3, L–O). Levels of this fusion protein drop somewhat during the first meiotic division (Figure 3, M and N) and become essentially undetectable before the end of the second meiotic division (Figure 3O). We conclude that Deg-A and Deg-B are sufficient in combination to trigger the rapid degradation of mNG::OMA-2 during meiosis, although this event is temporally delayed relative to GFP::LIN-41 (Figure 3P).

*oma-1* and *oma-2* share redundant functions during both oocyte and early embryo development (Detwiler *et al.* 2001; Guven-Ozkan *et al.* 2008). Double mutants carrying strong loss-of-function alleles (e.g., *oma-l(zu405te33); oma-2(te51))* are sterile with a defect in meiotic maturation (Detwiler *et al.* 2001; Table 1). For the most part, the embryonic functions of *oma-1/2* have been studied using conditions that reduce, but do not eliminate, OMA-1/2 function in embryos, such as double RNA interference (RNAi) or reduction-of-function alleles that are incompletely sterile (Nishi and Lin 2005; Guven-Ozkan *et al.* 2008). In *oma-I(zu405te33); oma-2(tn1764)* double mutants, OMA-2 is expressed during oogenesis but eliminated prematurely from embryos. Consequently, these double mutants are very fertile but produce progeny that die during embryogenesis (Table 2). Thus, as a novel allele that specifically reduces embryonic OMA-2, *oma-2(tn1764)* may be useful for studying the embryonic functions of *oma-1/2.* Our initial observations indicate that young *oma-I(zu405te33); oma-2(tn1764)* embryos exhibit cell division defects and ectopic cleavage furrows (Figure S8B); similar defects have been reported after *oma-l/2(RNAi)* depletion (Li *et al.* 2009).

When combined with *oma-l(zu405te33), oma-2(tn1764)* exhibits a strongerembryonic phenotype than either *oma-2(cp145)* or *oma-2(tn1760).* However, the severity of the *oma-I(te33zu405); oma-2(tn1760)* double mutant phenotype relative to *oma-l(te33zu405); oma-2(cp145)* was somewhat surprising (Table 2; 60 vs. 12% embryonic lethality). One possibility for the stronger embryonic phenotype might be the reduction in mNG::Deg-A::OMA-2 levels observed in *oma-2(tn1760)* pronuclear-stage embryos (Figure 3E and Figure S8A). We examined this more closely by crossing each mNG-tagged *oma-2* allele into an *emb-30(tn377ts)* mutant background, *emb-30* encodes a subunit of the Anaphase Promoting Complex (APC), and adult *emb-30(tn377ts)* hermaphrodites upshifted to restrictive temperature (25°C) produce 1-cell embryos that arrest in metaphase of the first meiotic division (Furuta *et al.* 2000). Arrest in meiotic metaphase does not prevent or delay the elimination of GFP::LIN-41, which is independent of APC function (Spike *et al.* 2014a). We observed that mNG::OMA-2 is turned over in arrested meiotic embryos, but could typically be seen in 4 embryos in the uterus of *emb-30(tn377ts); oma-2(cp145)* hermaphrodites after a 5-7 hour upshift to 25°C (Figure S8C). In contrast, both of the LIN-41 Deg domain-containing OMA-2 proteins appeared to be less stable under the same conditions. mNG::Deg-A, Deg-B::OMA-2 was seen in 0-1 mNG-positive embryos and appeared to be the least stable (Figure S8E), as expected from our previous analysis (Figure 3 and Figure S7). mNG::Deg-A::OMA-2 was seen in 2 mNG-positive embryos and therefore appeared to be of intermediate stability (Figure S8D). Thus, although Deg-A is not sufficient for rapid elimination, it likely reduces the stability of mNG::Deg-A::OMA-2 in meiotic embryos. The consequent reduction in protein levels could contribute to the stronger *oma-2(tn1760)* embryonic phenotype, although it also seems possible that insertion of LIN-41 Deg-A at the N-terminus of OMA-2 might perturb the nearby TAF-4 binding domain (Figure 3, A-C), which is critical for the function of OMA-2 in embryos (Guven-Ozkan *et al.* 2008).

### GFP::LIN-41 is eliminated from embryos by SCF^SEL-10^

Several different Skp, Cullin, F-box (SCF)-containing E3 ubiquitin ligase complexes promote protein degradation during meiosis and early embryogenesis in *C. elegans* (Peel *et al.* 2012; Du *et al.* 2015; Beard *et al.* 2016). We initially used RNAi to knock down the functions of each of the six cullins identified in the *C. elegans* genome (Kipreos *et al.* 1996; Nayak *et al.* 2002) to determine whether an SCF-type E3 ligase is involved in the elimination of GFP::LIN-41. In general, RNAi was initiated in *lin-41(tn1541)* hermaphrodites at the L4 larval stage and GFP::LIN-41 was examined in adults, two days after the initiation of the RNAi treatment at 22°C. Of the six cullins we tested, only the *cul-l(RNAi)-*treated animals produced multiple young embryos with faint GFP::LIN-41 (n=12), suggesting that CUL-1 may be required to eliminate GFP::LIN-41 from embryos. *rrf-l(pk1417)* mutants are RNAi-defective in certain somatic cells, including the somatic gonad, but are sensitive to RNAi in the germline (Sijen *et al.* 2001; Kumsta and Hansen 2012). Treatment of *rrf-l(pk1417) lin-41(tn1541)* hermaphrodites with *cul-1(RNAi)* also resulted in the failure to eliminate GFP::LIN-41 from early embryos (n=54; Figure 4C). Together, these results suggest that a germline-expressed CUL-1-containing SCF E3 ubiquitin ligase may eliminate GFP::LIN-41 from early embryos. At least three of the *C. elegans* cullins, *cul-1, cul-2* and *cul-3,* are important for normal embryonic development. We observed highly penetrant embryonic lethality after treating *rrf-l(pk1417) lin-41(tn1541)* and *lin-41(tn1541)* animals with *cul-l(RNAi)* and *cul-3(RNAi),* respectively. However, *cul-2(RNAi)* did not result in any obvious phenotypes. We therefore examined *lin-41(tn1541); cul-2(or209ts)* adults upshifted to 25°C as L4s to assess whether *cul-2* is important for the elimination of GFP::LIN-41 from embryos. GFP::LIN-41 was eliminated normally from the dead embryos produced by *cul-2(or209ts)* parents at restrictive temperature (n=71), suggesting that a CUL-2-containing SCF E3 ubiquitin ligase is not involved in the elimination of GFP::LIN-41 from embryos.

**FIGURE 4.**
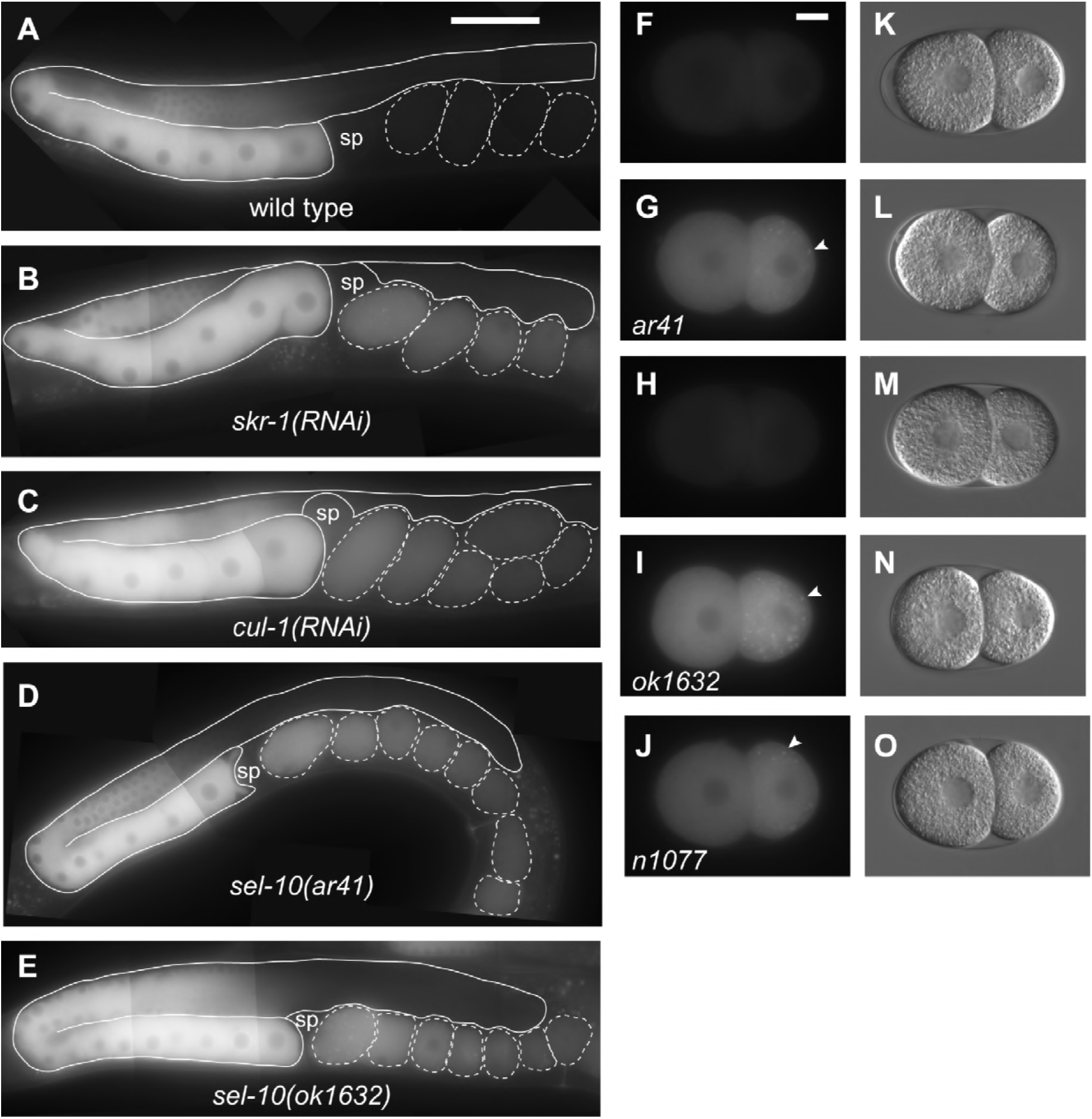
Subunits of the SCF^SEL-10^ E3 ubiquitin ligase are required for the elimination of GFP::LIN-41 from early embryos. (A–E) Composite images of GFP::LIN-41 in adult *rrf-l(pk1417) lin-41(tn1541)* hermaphrodites fed control RNAi bacteria (A), and adult hermaphrodites with reduced SCF^sel-10^ E3 ubiqitin ligase activity (B-E): *lin-41(tn1541); skr-l(RNAi)* (B), *rrf-l(pk1417) lin-41(tn1541); cul-l(RNAi)* (C), *lin-41(tn1541); Ion-3(e2175)sel-10(ar41)* (D), and *lin-41(tn1541); sel-10(ok1632)* (E). 100 ms GFP exposures, brightened slightly (and equivalently) to better visualize embryonic GFP::LIN-41; scale bar, 50 μm. (F-O) Images of 2-cell embryos removed from the uterus of hermaphrodites were imaged for GFP (F–J) and DIC (K-O); the genotypes were as follows: *lin-41(tn1541); lon-3(e2175)* (F, K), *lin-41(tn1541); lon-3(e2175) sel-10(ar41)* (G, L), *lin-41(tn1541)* (H, M), *lin-41(tn1541);sel-10(ok1632)* (l,N), and *lin-41(tn1541);sel-10(n1077)* (J, 0). Arrowheads indicate a few of the GFP::LIN-41 aggregates in the posterior blastomeres of *sel-10* mutant embryos, which likely correspond to P granules. 300 ms GFP exposures; scale bar, 10 μm.

In SCF-type E3 ligases, cullins interact with Skp-l-related proteins. Twenty-one Skpl-related *(skr)* genes have been identified in *C. elegans,* and RNAi experiments suggest the closely related *skr-1* and *skr-2* genes function in the germline and early embryo (Nayak *et al.* 2002; Yamanaka *et al.* 2002; Shirayama *et al.* 2006; Fox *et al.* 2011; Mohammad *et al.* 2018). In addition, both SKR-1 and SKR-2 can interact with CUL-1 (Nayak *et al.* 2002, Yamanaka *et al.* 2002). We therefore examined whether *skr-l(RNAi),* which likely reduces the function of both *skr-1* and *skr-2,* would prevent the elimination of GFP::LIN-41 from early embryos, *lin-41(tn1541); skr-l(RNAi)* animals produced embryos with defects in the elimination of GFP::LIN-41 two days after RNAi treatment (n=26; Figure 4B). Treatment of *rrf-l(pk1417) lin-41(tn1541)* animals with *skr-l(RNAi)* also prevented the elimination of GFP::LIN-41 from early embryos (n=14). In addition, the *rrf-l(pk1417) lin-41(tn1541)* mutants treated with *skr-l(RNAi)* for two days at 22°C exhibited defects in germline morphology and embryo production that are consistent with the phenotypes previously described after *skr-l/2(RNAi)* (Nayak *et al.* 2002).

At least three F-box-containing substrate recognition subunits, LIN-23, PROM-1, and SEL-10, are thought to function with either SKR-1 or SKR-2 and CUL-1 in the *C. elegans* germline or early embryos (Peel *et al.* 2012; Du *et al.* 2015; Kisielnicka *et al*. 2018; Mohammad *et al.* 2018). LIN-23, SEL-10 and theirvertebrate orthologs (β-TrCP and Fbw7, respectively) regulate the cell cycle and cell cycle-dependent protein degradation (Kipreos *et al.* 2000; Nakayama and Nakayama 2005; Welcker and Clurman 2008; de la Cova and Greenwald 2012). Because the rapid elimination of GFP::LIN-41 appears to be coupled to meiotic maturation, a cell-cycle event, we used RNAi to knock down the activities of *lin-23* and *sel-10* as a first step toward the analysis of candidate F-box proteins. *lin-23(RNAi)* had no effect on the elimination of GFP::LIN-41 from *rrf-1(pk1417) lin-41(tn1541)* embryos (n=52; Figure S9C). Consistent with this observation, mutations designed to prevent the phosphorylation of a possible β-TrCP binding site near the amino-terminus of LIN-41 (amino acids 32-38) also do not prevent the elimination of GFP::LIN-41 from embryos *lin-41(tn1541tn1668);* Figure S5, D and E). However, *sel-10(RNAi)* did prevent the elimination of GFP::LIN-41 from *rrf-1(pk1417) lin-41(tn1541)* embryos (n=17). Similarly, the elimination of GFP::LIN-41 from young embryos is prevented by the strong loss-of-function mutations *sel-10(ok1632)* and *sel-10(ar41)* (Figure 4, D, E, G, and I). All of our sel-*10(ar41)* strains also contain *lon-3(e2175),* a convenient cis-linked marker that encodes a cuticle collagen (Nyström *et al*. 2002, Suzuki *et al.* 2002). GFP::LIN-41 is eliminated normally from *lon-3(e2175)* mutant embryos (Figure 4F). Finally, we observed that *sel-10(n1077),* which has both gain-of-function and loss-of-function properties (Jäger *et al.* 2004), fails to eliminate GFP::LIN-41 from early embryos (Figure 4J). Genetic and physical interactions indicate that SEL-10 and SKR-1 function together in *C. elegans* (Killian *et al.* 2008; Kisielnicka *et al.* 2018). Because our *skr-1(RNAi)* experiments are likely to also target *skr-2* (Nayak *et al.* 2002), we are unable to parse out the relative roles of SKR-1 and SKR-2 at this time. Collectively, these observations suggest that a germline-expressed SCF^SEL-10^ E3 ubiquitin ligase containing SKR-1/2, CUL-1 and SEL-10 is likely involved in the elimination of GFP::LIN-41 from early embryos (Figure 5E).

**FIGURE 5.**
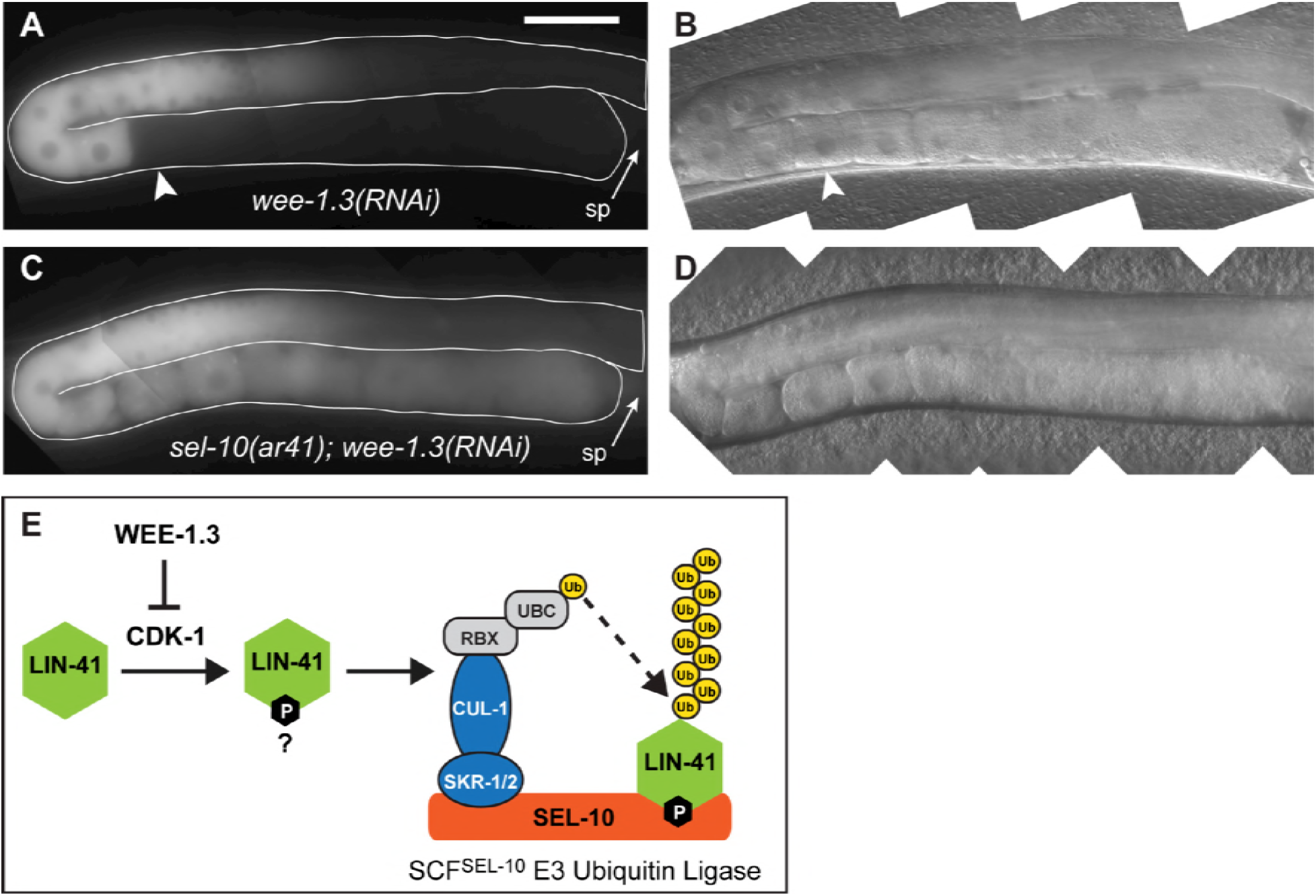
SEL-10 is required for the WEE-1.3-inhibited degradation of GFP::LIN-41. (A–D) Composite GFP (A, C) and DIC (B, D) images of *lin-41(tn1541); lon-3(e2175); wee-1.3(RNAi)* (A, B) and *lin-41(tn1541); Ion-3(e2175)sel-10(ar41); wee-1.3(RNAi)* (C, D) animals. GFP::LIN-41 is prematurely eliminated from oocytes by *wee-1.3(RNAi)* (arrowhead), but persists in abnormal oocytes nearthe spermatheca (sp, arrow) in *sel-10(ar41); wee-1.3(RNAi)* animals (C, D), suggesting that SEL-10 is required for this process. 150 ms GFP exposures, brightened slightly; scale bar, 50 μm. (E) A simple model for the elimination of LIN-41 (green) that incorporates the known molecular functions of WEE-1.3 kinase, cyclin-dependent kinase (CDK-1) and subunits of the SCF^SEL-10^ E3 ubiquitin ligase. In brief, we hypothesize that SEL-10 (orange) may recognize phosphorylated LIN-41 (green) and trigger its ubiquitin-mediated degradation in collaboration with the other SCF E3 ubiquitin ligase subunits, SKR-1/2 (blue) and CUL-1 (blue). CUL-1 orthologs bind RING finger proteins (RBX, gray), which recruit a ubiquitin-conjugating enzyme (UBC, gray) that catalyzes the transfer of ubiquitin (yellow) to protein substrates, such as LIN-41. Subsequent recruitment of poly-ubiquitinated substrates to the proteasome results in degradation (not shown). This model is consistent with the epistatic relationship between *wee-1.3(RNAi)* and *sel-10(ar41)* with respect to the elimination of GFP::LIN-41, but other models are also possible.

### SEL-10 functions through LIN-41 degradation domains

LIN-41 can be detected in *sel-10(ok1632)* mutant but not wild-type embryos by western blot analysis (Figure S6C), indicating that endogenous and GFP-tagged LIN-41 behave similarly. We hypothesized that the Deg domains are likely used to target LIN-41 for degradation by SCF^SEL-10^. To test this hypothesis, we examined whether the premature elimination of mNG::Deg-A, Deg-B::OMA-2, which is mediated by the LIN-41 Deg domains, is prevented in *sel-10* mutant embryos. Although mNG::Deg-A, Deg-B::OMA-2 is eliminated by the pronuclear stage in otherwise wild-type 1-cell embryos, mNG::Deg-A, Deg-B::OMA-2 levels remain high in *lon-3(e2175) sel-10(ar41)* embryos at the same stage of embryonic development (Figure 3, F, G, J, and K, and Figure S7, G, H, K, and L). As for GFP::LIN-41, this is not caused by the cis-linked marker *lon-3(e2175)* (Figure S7, I and J). These observations suggest that *sel-10(ar41)* should suppress the completely penetrant maternal-effect lethal phenotype exhibited by *oma-I(zu405te33); oma-2(tn1764)* mutants (Table 2). Consistent with this expectation, *oma-I(zu405te33); oma-2(tn1764) lon-3(e2175) sel-10(ar41)* animals produce hatchlings and can be maintained as a homozygous strain (Table 2). However, we note that *sel-10(ar41)* is a relatively weaksuppressorofthe *oma-l(zu405te33); oma-2(tn1764)* maternal-effect lethal mutant phenotype, since only 10-15% of the embryos produced by *oma-l(zu405te33); oma-2(tn1764) lon-3(e2175) sel-10(ar41)* animals hatch. This observation is consistent with the possibility that mNG::Deg-A, Deg-B::0MA-2 may only be partially functional in the 1-cell embryo. One possibility is that the Deg-A, Deg-B insertion perturbs OMA-2 function. Additionally, *sel-10(ar41)* mildly reduces mNG::OMA-2 accumulation in the germline likely through effects on GLD-1 (see below). Although the degradation of OMA-1, and presumably OMA-2, appears to be mediated by several SCF E3 ubiquitin ligases, SEL-10 has not been implicated in this process (Shirayama *et al.* 2006; Du *et al.* 2015). Indeed, mNG::OMA-2 is degraded at the expected time in *oma-2(cp145) lon-3(e2175) sel-10(ar41)* embryos (Figure S7, C and D). Likewise, mNG::Deg-A, Deg-B::OMA-2 levels only remain high until the end of the 1-cell stage in *oma-2(tn1764) lon-3(e2175) sel-10(ar41)* embryos, when the degradation of OMA-2 is normally initiated (Figure S7, K and L). We conclude that SEL-10 is not required for the elimination of OMA-2 and likely functions through the LIN-41 Deg domains to promote the proteolytic degradation of LIN-41 and mNG::Deg-A, Deg-B::OMA-2 during meiosis.

### SEL-10 is required for the CDK-1-dependent elimination of GFP::LIN-41

Substrate recognition subunits such as SEL-10 recognize their targets by binding to short linear sequence motifs called degrons (Lucas and Ciulli 2017). LIN-41 Deg domains were therefore examined for sequences similar to the SEL-10/Fbw7/Cdc4 degron consensus sequence ΦΦ[pT/pS]PXX[pT/pS/E/D], where Φ represents a hydrophobic amino acid. This degron is commonly referred to as a Cdc4 phosphodegron or CPD; it contains two essential residues, a phosphorylated residue that is typically a phosphothreonine, immediately followed by a proline (pTP) (Nash *et al.* 2001). Residues surrounding LIN-41 T83, which is important for the elimination of LIN-41 from embryos, are poor matches to this consensus sequence (sequence FDTPPSM, mismatches are underlined; Figure S10). The best match to a high-affinity CPD appears to be around residue T340 in the LIN-41 Deg-B2 domain (sequence LATPMSS; Figure S10). This was the only candidate Fbw7 binding site identified in LIN-41 using the Eukaryotic Linear Motif (ELM) database (Gouw *et al.* 2018), which requires a perfect match to a relatively stringent consensus sequence. However, changing T340 to an alanine (T340A) (e.g.: *lin-41(tn1541tn1775)* has no effect on the elimination of GFP::LIN-41 from embryos (Figure 2C and Figure S4, K and L). Therefore, if SEL-10 binds directly to LIN-41 Deg domains it might recognize imperfect or lower-affinity degrons. We note that the SEL-10 ortholog Cdc4p utilizes multiple imperfect degrons to target the cell division protein Sic1p for degradation (Nash *et al.* 2001). Likewise, multiple weak degrons in an intrinsically disordered region of the c-Jun protein synergize to promote a high-affinity interaction with the SEL-10 ortholog Fbw7 (Csizmok *et al.* 2018). It seems plausible that SEL-10 might function similarly. Alternatively, the failure to eliminate LIN-41 from embryos could be an indirect consequence of the lack of SCF^SEL-10^. To begin to address this possibility, we sought to clarify the epistatic relationships between *sel-10* and other factors involved in the elimination of GFP::LIN-41 from embryos.

CDK-1 was previously shown to be required for the elimination of GFP::LIN-41 (Spike *et al.* 2014a). Likewise, *cdk-l(RNAi)* on *rrf-l(pk1417) lin-41(tn1541)* hermaphrodites prevents the elimination of GFP::LIN-41 from embryos (n=67; Figure S9, A and B). Therefore, germline-expressed CDK-1 likely promotes the elimination of GFP::LIN-41. CDK-1 is a conserved and essential cell-cycle regulator required for M-phase entry and progression during both meiotic and mitotic cell divisions (Boxem *et al.* 1999). Consequently, most *cdk-1* alleles are sterile, precluding the examination of GFP::LIN-41 in *cdk-1* mutant embryos. Two temperature-sensitive alleles of *cdk-1* that produce oocytes have been described; both cause a later embryonic arrest phenotype than the 1-cell meiotic arrest phenotype seen after *cdk-1(RNAi)* (Boxem *et al.* 1999; Shirayama *et al.* 2006). Furthermore, although both mutations alter residues in the T loop/activation domain of CDK-1, neither *cdk-1(ts)* allele causes obvious cell-cycle defects (Shirayama *et al.* 2006). We examined GFP::LIN-41 in *cdk-l(ne2257ts)* animalsat the restrictive temperature and found that GFP::LIN-41 disappears normally from embryos (n=57; Figure S9, E and F). Similarly, GFP::LIN-41 disappears normally in *cks-1(ne549ts)* mutant embryos (n=33; Figure S9, G and H), which phenotypically resemble *cdk-l(ne2257ts)* embryos at the restrictive temperature (Shirayama *et al.* 2006). Thus, the subset of CDK-1 activities affected by *cdk-1(ne2257ts)* does not include either the elimination of GFP::LIN-41 or entry into meiotic M phase.

Kinases other than CDK-1 might play a role in the SEL-10-mediated elimination of LIN-41. Indeed, WEE-1.3, a kinase that negatively regulates CDK-1 (Burrows *et al.* 2006), prevents the premature elimination of GFP::LIN-41 from oocytes (Spike *et al.* 2014a). However, our attempts to identify additional kinases that affect the elimination of GFP::LIN-41 have so far been unsuccessful. For example, the mitogen-activated protein (MAP) kinase MPK-1 is active in the late stages of oogenesis and is an important regulator of oocyte meiotic maturation (Lee *et al.* 2007). Furthermore, as a proline-directed serine/threonine kinase, MPK-1 could potentially phosphorylate CPDs in LIN-41 Deg domains. However, GFP::LIN-41 disappears normally in *mpk-1(galllts)* embryos at the restrictive temperature (n=97; Figure S9, I-L). Likewise, the Aurora kinase AIR-2 is present and active in maturing oocytes (Schumacher *et al.* 1998), but GFP::LIN-41 is eliminated normally after *air-2* gene function is attenuated by *air-2(or207ts)* (n=14), by *air-2(RNAi)* (n=40), or after *air-2(RNAi)* on *air-2(or207ts)* mutants (n=38; Figure S9, M and N). Other single kinase knock-down or elimination experiments that have failed to affect the elimination of GFP::LIN-41 from *lin-41(tn1541)* embryos include *gsk-3(RNAi)* (n=32), *cdk-2(RNAi)* (n=24), *plk-l(RNAi),* and *mbk-2(pk1427)* (Spike 2014a). Since LIN-41 functions to inhibit CDK-1 activation for M-phase entry during meiotic maturation (Spike *et al.* 2014a), CDK-1 may be the chief effector kinase mediating feedback regulation of wild-type LIN-41.

SEL-10/Fbw7/Cdc4p degrons are only activated after being phosphorylated by a proline-directed serine/threonine kinase such as CDK-1. Thus, prior phosphorylation by CDK-1 might be required for SEL-10 to directly interact with degrons in the LIN-41 Deg domains. *cdk-l(RNAi)* and *sel-10(lf)* cause the same phenotype with respect to GFP::LIN-41 degradation, precluding a direct analysis of their epistatic relationship. However, it is possible to examine this relationship indirectly through *wee-1.3* (Burrows *et al.* 2006). GFP::LIN-41 is eliminated prematurely when *wee-1.3* function is attenuated by RNAi; this occurs in both wild-type (Spike *et al.* 2014a) and *lon-3(e2175)* mutants (n=21; Figure 5, A and B). When the same experiment is performed in *lon-3(e2175) sel-10(ar41)* mutants, however, GFP::LIN-41 is not eliminated prematurely. Instead, GFP::LIN-41 persists, typically at reduced levels, in the proximal oocytes of *lon-3(e2175) sel-10(ar41); wee-1.3(RNAi)* animals (n=51; Figure 5, C and D). Because CDK-1 is prematurely activated, *wee-1.3(RNAi)* oocytes mature prematurely and exhibit numerous defects (Burrows *et al.* 2006). Obvious oocyte abnormalities caused by strong *wee-1.3(RNAi)* are evident in *sel-10(ar41)* mutants, confirming that these animals are competent to respond to *wee-1.3(RNAi)* (Figure 5, compare B and D). We conclude that the epistatic relationship between *wee-1.3* and *sel-10* is consistent with the model shown in Figure 5E, which postulates that active CDK-1 promotes the phosphorylation of LIN-41 and its subsequent destruction by an Scf^SEL-10^ E3 ubiquitin ligase.

### Embryonic LIN-41 does not strongly inhibit the expression of mRNAs repressed by LIN-41

LIN-41 represses the translation of several different mRNAs during oogenesis (Spike *et al.* 2014b; Tsukamoto *et al.* 2017). Their protein products normally begin to accumulate in late oogenesis or early embryogenesis, and some are essential for normal development (Gomes *et al.* 2001; Leacock and Reinke 2008; Tsukamoto *et al.* 2017). We therefore anticipated that the failure to eliminate LIN-41 would result in the ectopic repression of these mRNAs, and that this, in turn, might result in embryo or oocyte abnormalities. However, the *lin-41(tn1767)* mutant, which fails to eliminate LIN-41[T83A] from early embryos, appears essentially wild-type at 20°C (Table 1). Similarly, *sel-10(ok1632)* and *sel-10(ar41)* mutants, which fail to eliminate LIN-41 from early embryos, produce large broods of progeny at 20°C that are similar in size to genotype-matched controls (Table 3). Indeed, we only observed a moderate decrease in fertility when *sel-10(ok1632)* mutants were grown at an elevated temperature (25°C; Table 3). We therefore decided to examine the amount of protein made by several LIN-41 target mRNAs *(spn-4, meg-1,* and *orc-1* mRNAs, respectively) in strains that fail to eliminate LIN-41 from embryos, as this should provide a sensitive way to monitor LIN-41 translational repression activity. Protein expression was examined using fluorescently-tagged alleles of each gene; the proteins made by *spn-4(tn1699[spn-4::gfp::3xflag]), meg-1(tn1724[gfp::3xflag::meg-1])* and *orc-1(tn1732[mng::3xflag::orc-1)* were previously shown to be ectopically or prematurely expressed in *lin-41(lf)* oocytes (Tsukamoto *et al.* 2017). As described below, only minor differences in protein expression were observed in *lin-41(tn1767)* and *sel-10(ar41)* embryos and oocytes (Figure 6 and Figure S11). Collectively, these observations suggest that the ectopic LIN-41 present in *lin-41(tn1767)* and *sel-10(lf)* embryos is largely ineffective at repressing translation.

**Table 3.** *sel-10* mutant brood sizes at 20° and 25°C

| GENOTYPE | T (°C) | BROOD SIZE |
| --- | --- | --- |
| wild type | 20 | 304.0 ± 31.1 (n=29) |
| wild type<sup>a</sup> | 25 | 266.8 ± 39.0 (n=19) |
| *sel-10(ok1632)*<sup>a</sup> | 20 | 258.3 ± 67.7 (n=30) |
| *sel-10(ok1632)*<sup>a</sup> | 25 | 72.2 ± 34.0<sup>b</sup> (n=30) |
| *lin-41(tn1487ts); sel-10(ok1632)*<sup>c</sup> | 20 | 3.2 ± 3.4 (n=53) |
| *lin-41(tn1487ts)*<sup>d</sup> | 20 | 41.4 ± 23.8 (n=36) |
| *lon-3(e2175)* | 20 | 294.5 ± 36.7 (n=20) |
| *lon-3(e2175) sel-10(ar41)* | 20 | 280.2 ± 40.1 (n=20) |

<sup>a</sup> Newly fertilized embryos were collected at 15°C and shifted to 25°C.
<sup>b</sup> Approximately 10.0 ± 5.3% of *sel-10(ok1632)* hermaphrodites (n=3568) are infertile, exhibiting incompletely penetrant sterility or maternal-effect lethality when grown and examined for seven generations at 25°C.
<sup>c</sup> The progeny of *sel-10(ok1632); lin-41(tn1487ts)/hT2[qIs48]* hermaphrodites.
<sup>d</sup> The progeny of *lin-41(tn1487ts)/hT2(qIs48)* hermaphrodites.

LIN-41 mediates 3’-UTR-dependent translational repression of *spn-4,* and *spn-4* mRNA is the most abundant and enriched mRNA in LIN-41 RNPs (Tsukamoto *et al.* 2017). SPN-4::GFP is faint, but visible, inlor2 proximal oocytes and rapidly accumulates during the oocyte-to-embryo transition (Tsukamoto *et al.* 2017). This pattern, and the amount of SPN-4::GFP in early embryos, is largely unaffected by *lin-41(tn1767)* and *sel-10(ar41)* at 20°C (Figure 6, A, B, G, and H, and Figure S11, A, B, K, and L). Quantification of GFP levels revealed no differences in SPN-4::GFP levels in *lin-41(tn1767); spn-4(tn1699)* 1- and 2-cell embryos and a slight reduction in SPN-4::GFP levels in *spn-4(tn1699) lon-3(e2175) sel-10(ar41)* 2-cell embryos relative to age and genotype-matched controls (Figure 6, K and L). In these quantitative experiments, we analyzed the anterior cytoplasm of 1- and 2-cell embryos and did not include the bright puncta of SPN-4::GFP evident in the posterior cytoplasm (see Materials and Methods). Finally, we also failed to identify any apparent differences in SPN-4::GFP accumulation or intensity in *spn-4(tn1699) lon-3(e2175) sel-10(ar41)* (n=26) and *spn-4(tn1699) lon-3(e2175)* (n=19) animals upshifted as L4s to 25°C.

**FIGURE 6.**
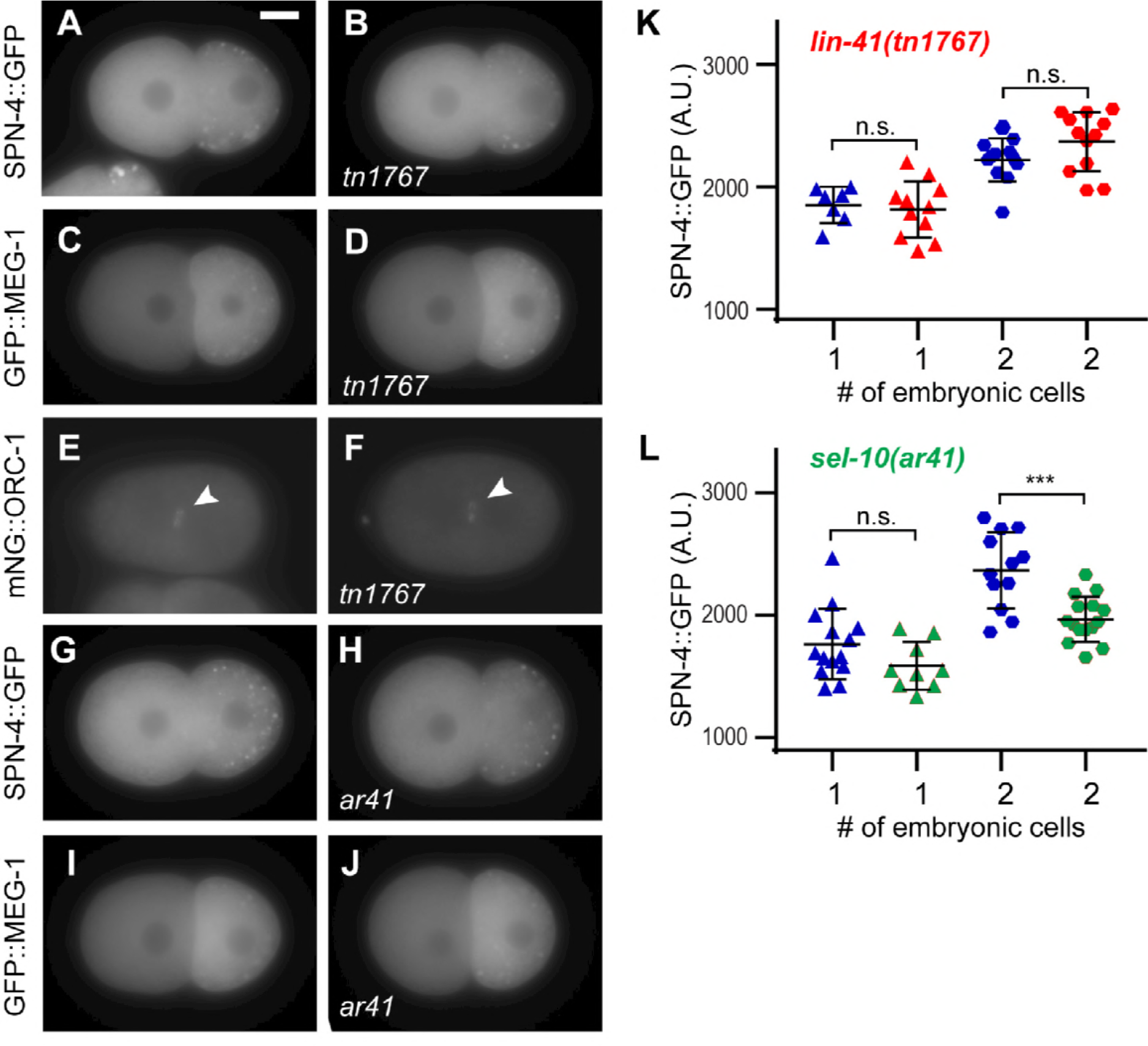
Persisting LIN-41 or LIN-41[T83A] does not strongly inhibit the expression of LIN-41 targets of translational repression in young embryos. (A–J) Young embryos express similar levels ofSPN-4::GFP (A, B, G, and H), GFP::MEG-1 (C, D, I, and J) and mNG::ORC-1 (arrowhead in E, F) when ectopic LIN-41[T83A] (B, D, and F; *lin-41(tn1767)* mutant embryos), ectopic LIN-41 (H, J; *sel-10(ar41)* mutant embryos) or normal (undectable) levels of LIN-41 (B, D, F, H, and J) are present. Exposures were 100 ms for SPN-4::GFP, 200 ms for GFP::MEG-1 and 600 ms for mNG::ORC-1; scale bar, 10 μm. (K) Quantification of the intensity of SPN-4::GFP expression in *spn-4(tn1699)* and *Hn-41(tn1767);spn-4(tn1699)* 1 and 2-cell embryos. No significant differences were seen (n.s.). (L) Quantification of the intensity of SPN-4::GFP expression in *spn-4(tn1699); lon-3(e2175)* and *spn-4(tn1699); lon-3(e2175) sel-10(ar41)* 1 and 2-cell embryos. Levels appeared to be slightly lower in the *sel-10(ar41)* 2-cell embryos (P<.001). Note that the slightly reduced level of SPN-4::GFP in image (H) relative to image (G) accurately illustrates the very modest magnitude of this difference in expression.

GFP::MEG-1 expression is evident somewhat earlier in oogenesis than SPN-4::GFP and seems to accumulate more slowly. Again, the pattern and amount of GFP::MEG-1 was largely unaffected in *lin-41(tn1767)* and *sel-10(ar41)* mutants at 20°C (Figure 6, C, D, I, and J, and Figure S11, C, D, G, and H). GFP::MEG-1 has a complex pattern of accumulation in the early embryo; it appears to be eliminated from somatic blastomeres and localizes, at least partially, to P granules (Figure 6C and Figure S11C), similarto what was previously described by Leacock and Reinke (2008) for endogenous MEG-1. Due to these complexities, we quantified GFP::MEG-1 levels in the cytoplasm of proximal oocytes instead of embryos. There were no differences in GFP::MEG-1 levels in *lin-41(tn1767); meg-1(tn1724)* animals and only a slight reduction in the – 1 oocytes of *lon-3(e2175) sel-10(ar41); meg-1(tn1724)* animals relative to controls. Finally, we examined *lon-3(e2175)sel-10(ar41); meg-1(tn1724)* (n=17) and *lon-3(e2175); meg-1(tn1724)* (n=15) animals upshifted as L4s to 25°C, but again failed to identify any apparent differences in GFP::MEG-1 accumulation.

mNG::ORC-1 is not visibly expressed in oocytes but becomes increasingly evident in embryos as they develop (Tsukamoto *et al.* 2017). mNG::ORC-1 associates with chromatin at certain stages of the cell cycle (Sonneville *et al.* 2012), and is faintly visible in 1-cell embryos during metaphase of the first mitotic division (Figure 6E). mNG::ORC-1 was only examined in *lin-41(tn1767)* mutants at 20°C. As for SPN-4::GFP and GFP::MEG-1, the pattern and amount of mNG::ORC-1 was largely unaffected by *lin-41(tn1767)* (Figure 6, E and F, and Figure S11, E and F). Most importantly, the small amount of mNG::ORC-1 visible in 1-cell embryos was not obviously reduced in the *lin-41(tn1767)* background. Because LIN-41 is a potent translational repressor of *spn-4, meg-1,* and *orc-1* (Tsukamoto *et al.* 2017), we conclude that a mechanism distinct from SCF^SEL10^-mediated degradation antagonizes LIN-41 function to promote their expression during the late stages of oogenesis and the OET.

Because moleculartests failed to reveal an increase in LIN-41 activity in *sel-10* mutants, we also tested the ability of a *sel-10(ok1632)* strong loss-of-function mutation to suppress the temperature-sensitive *lin-41(tn1487ts)* allele at a semi-permissive temperature. This was found not to be the case; rather, *sel-10(ok1632)* enhanced the *lin-41(tn1487ts)* defects (Table 3). Taken together, these results indicate that the regulation of LIN-41 by *sel-10* is nonessential.

### SEL-10 promotes the elimination of GLD-1 from oocytes

GLD-1 is a translational repressor that, like LIN-41, controls and coordinates oocyte differentiation and cell cycle progression (Francis *et al.* 1995a,b; Jones *et al.* 1996). In *gld-1(q485)* null mutants, pachytene-stage oocytes re-enter the mitotic cell cycle and form a tumor (Francis *et al.* 1995a,b). GLD-1 also has redundant functions to inhibit the proliferative fate of germline progenitor cells and to promote their entry into the meiotic pathway of development during oogenesis and spermatogenesis, as well as a function to promote spermatogenesis in hermaphrodites (Francis *et al.* 1995a,b; Kadyk and Kimble 1998). GLD-1 is abundantly expressed during the early and middle stages of meiotic prophase, but eliminated from oocytes as they progress from late pachytene through diplotene and to diakinesis during the later stages of oocyte development (Jones *et al.* 1996). GLD-1 binds to, and represses the translation of, many mRNAs that are normally translated in oocytes (Lee and Schedl 2001; Lee and Schedl 2004; Schumacher *et al. 2005;* Wright *et al.* 2011; Jungkamp *et al.* 2011; Scheckel *et al.* 2012). Thus, it has generally been assumed that the elimination of GLD-1 from oocytes permits the translation of these mRNAs (reviewed in Lee and Schedl 2010). Although it occurs at an earlier stage of oocyte development, this model is analogous to what we originally hypothesized with respect to LIN-41. However, because the LIN-41 ectopically found in *sel-10* loss-of-function embryos appears to be insufficient to sustain translational repression, it seems likely that the activity of LIN-41 is also regulated by a non-proteolytic mechanism. Given the similarities between LIN-41 and GLD-1, we wondered whether GLD-1 might also be regulated by both proteolytic and non-proteolytic mechanisms.

To begin to approach this question, we investigated whether the elimination of GLD-1, like LIN-41, requires SEL-10. Surprisingly, we found that GLD-1::GFP and GLD-1 do indeed persist at elevated levels in the oocytes of *sel-10(ar41)* and *sel-10(ok1632)* mutants (Figure 7, A and B, and Figure S12, A, B, D, and E), indicating that LIN-41 and GLD-1 may be regulated similarly. Indeed, while we were completing this work, the failure to eliminate GLD-1 in a timely fashion from *sel-10(ok1632)* mutant oocytes was independently discovered by Kisielnicka *et al.* (2018). Their results suggest that both GLD-1 and the cytoplasmic polyadenylation element-xbinding protein CPB-3 are likely degraded by essentially the same SCF^SEL-10^ E3 ubiquitin ligase (Kisielnicka *et al.* 2018) that regulates LIN-41 (this work). Consistent with this hypothesis, they observed that slow-migrating isoforms of GLD-1, which are likely phosphorylated (Jeong *et al.* 2011), accumulate in *sel-10(ok1632)* mutants. In agreement with this finding, we also observe an increase in the slow-migrating isoforms of GLD-1 in both *sel-10(ok1632)* and *sel-10(ar41)* mutants relative to controls (Figure 7C).

**FIGURE 7.**
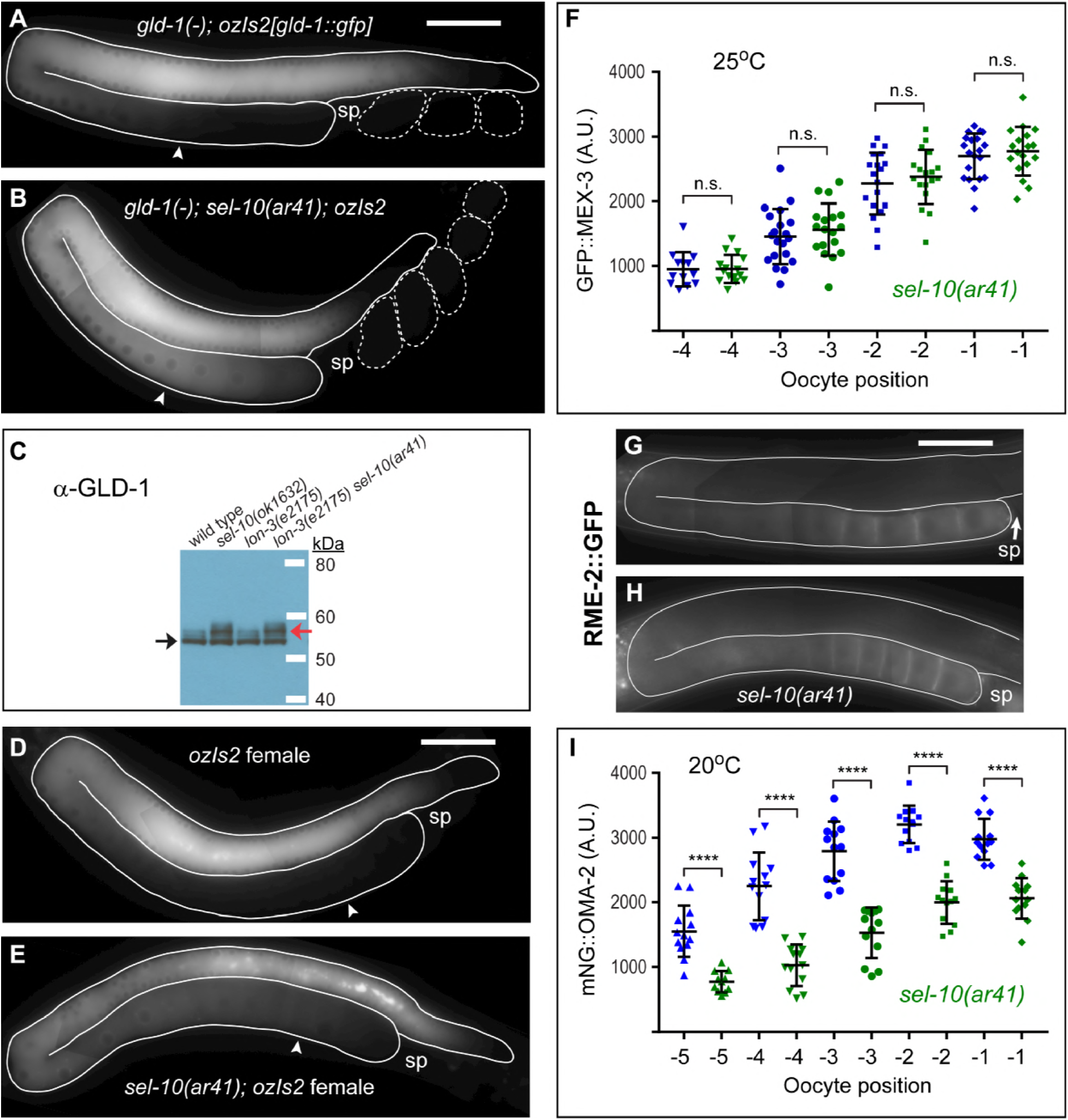
GLD-1 persists at elevated levels in the oocytes of *sel-10(ar41)* mutants. (A, B) Composite images of GLD-1::GFP in *gld-1(q485); lon-3(e2175); ozis2[gld-1::gfp]* (A) and *gld-1(q485); Ion-3(e2175)sel-10(ar41); ozis2[gld-1::gfp]* (B) adult hermaphrodites. GLD-1::GFP levels remain elevated in the proximal oocytes (e.g.: −4 oocytes, arrowheads) of *sel-10(ar41)* animals (B) relative to controls (A). 17 ms GFP exposures, brightened slightly. (C) Slow-migrating forms of GLD-1 (red arrow) are more abundant in *sel-lθ(lf)* adult hermaphrodites than in *sel-10(+)* controls, where the fast-migrating form of GLD-1 (black arrow) predominates. (D, E) Composite images of GLD-1::GFP in *fog-3(q470); lon-3(e2175); ozis2[gld-1::gfp]* (D) and *fog-3(q470); lon-3(e2175) sel-10(ar41); ozis2[gld-1::gfp]* females (E). GLD-1::GFP levels are elevated in the proximal oocytes (e.g.: −4 oocytes, arrowheads) of *sel-10(ar41)* females (B) relative to controls (A), although this is not as dramatic as in hermaphrodites. A somewhat longer GFP exposure (35 ms, brightened slightly) was needed than in (A and B), likely due to the presence of endogenous GLD-1. (F) Quantification of the intensity of GFP::MEX-3 in the proximal oocytes of *lon-3(e2175); mex-3(tn1753)* and *Ion-3(e2175)sel-10(ar41); mex-3(tn1753)* hermaphrodites at 25°C. No significant differences were seen (n.s.). (G-H) Composite images of *lon-3(e2175); pwis116[rme-2p::rme-2::GFP::rme-2 3’UTR]* (G) and *lon-3(e2175) sel-10(ar41); pwlsll? [rme-2p::rme-2::GFP::rme-2 3’UTR]* (H) hermaphrodites at 22°C. 300 ms GFP exposures. Neither target of GLD-1 translational repression (MEX-3, RME-2) was strongly or even marginally reduced in expression in *sel-10(ar41)* oocytes. (I) Quantification of the intensity of mNG::OMA-2 in the proximal oocytes of *oma-2(cp145) lon-3(e2175)* and *oma-2(cp145) lon-3(e2175) sel-10(ar41)* hermaphrodites at 20°C. Differences in expression were highly significant (P<.0001, indicated by 4 asterisks), but relatively modest in magnitude. For example, we measured a 37% reduction in average fluorescence in the −2 oocytes of *sel-10(ar41)* animals relative to the same oocytes in control animals. Scale bar, 50 μm (A, B, D, E, G, and H).

It was recently proposed that sperm trigger the proteasome-dependent elimination of GLD-1 from oocytes such that a GFP::GLD-1 transgene (an N-terminal fusion) was expressed at higher levels in unmated females than in mated females or hermaphrodites (Bohnert and Kenyon 2017). We therefore decided to examine the localization of a rescuing GLD-1::GFP transgene (a C-terminal fusion) in both wild-type and *sel-10(ar41)* mutant females, which lack sperm. However, in our experiments, GLD-1::GFP did not persist at elevated levels in female oocytes; instead, it was eliminated from oocytes in both the presence and absence of sperm (Figure 7, A and D, Figure S12, A and C). Likewise, endogenous GLD-1, detected with specific antibodies (Jan *et al.* 1999), also disappeared from oocytes in both hermaphrodites and females (Figure S12, D and F). However, GLD-1::GFP levels remained elevated in the oocytes of *sel-10(ar41)* mutant females (Figure 7E). Oocytes remain in the gonad for an extended period of time in the absence of sperm (McCarter *et al.* 1999). Indeed, we noticed that there seemed to be relatively less GLD-1::GFP in the *sel-10(ar41)* oocytes offemales as compared to hermaphrodites, possibly as a result of sel-10-independent protein turnover. From these results, we conclude that the sel-10-dependent elimination of GLD-1::GFP is sperm-independent. Furthermore, the expression patterns of GLD-1::GFP, which rescues the *gld-1(q485)* null mutation to fertility (Schumacher *et al.* 2005; Figure 7A), and endogenous GLD-1 (Figure S12, D and F), fail to support the hypothesis that sperm trigger the elimination of GLD-1 from oocytes. We have tested the GFP::GLD-1 transgene *(axis1498[pie-1p::gfp::gld-1::gld-1 3’UTR, unc-119(+)]¦* Merritt *et al.* 2008) used by Bohnert and Kenyon (2017) to monitor GLD-1 expression in females; however, we found that it fails to rescue *gld-1(q485)* null mutants to fertility. Adult *gld-1(q485); axis1498* hermaphrodites are invariably sterile and exhibit a range of phenotypes from a tumorous phenotype that is equivalent to the null allele to the production of abnormal oocytes. We analyzed 192 progeny from *gld-1(q485)/+; axis1498* adult hermaphrodites; 53 (27.6%) were *gld-1(q485); axis1498* and were sterile, consistent with the inability of *axis1498* to provide wild-type *gld-1* function. We conclude that the increased expression of GFP::GLD-1 observed in the oocytes of *axis1498* females (Bohnert and Kenyon 2017) is most likely a transgene expression artifact and does not reflect the expression and regulation of endogenous GLD-1.

### Ectopic GLD-1 in proximal oocytes does not strongly inhibit the expression of mRNAs repressed by GLD-1

As for LIN-41, we examined whether the mRNA targets of GLD-1-mediated translational repression are ectopically repressed in *sel-10(ar41)* mutant oocytes. Studies of GLD-1 function in the proliferative versus meiotic entry decision of germline progenitor cells demonstrate that GLP-l/Notch signaling functions to inhibit GLD-1 accumulation in the distal end of the germline. When GLD-1 accumulates ectopically in *glp-1* mutants, or double mutants affecting the Pumilio and FBF proteins FBF-1 and FBF-2, germline progenitor cells fail to proliferate and prematurely enter the meiotic pathway of development (Crittenden *et al.* 2002; Hansen *et al.* 2004). Thus, our initial expectation was that ectopic GLD-1 expression in proximal oocytes in strong *sel-10* loss-of-function mutants might exert substantial effects on the repression of its mRNA targets. As for LIN-41, this proved not to be the case; only subtle or modest effects were observed as described below.

GLD-1 binds the 3’-UTR of the *spn-4* mRNA (Jungkamp *et al.* 2011), which we initially examined as a LIN-41 target, and GLD-1 appears to repress SPN-4 accumulation in the distal germline (Mootz *et al.* 2004). As described previously, SPN-4::GFP expression was not strongly affected by the *sel-10(ar41)* mutation (Figure S11, I-L) despite the ectopic expression of both GLD-1 and LIN-41 (Figure 4 and Figure 7). MEX-3 is expressed in proximal oocytes and also appears to be repressed by GLD-1 (Mootz *et al.* 2004; Jungkamp *et al.* 2011). We used the fluorescently-tagged *mex-3(tn1753[gfp::3xflag::mex-3])* allele to quantitatively examine the expression of GFP::MEX-3 in these oocytes at both 20° and 25°C. GFP::MEX-3 levels were not reduced in *sel-10(ar41)* oocytes at either temperature, but were generally very similar to the wild-type controls (Figure 7F, and Figure S12H). In addition, we examined the expression of the yolk receptor RME-2 (Grant and Hirsh 1999), a well-established target of GLD-1-mediated translational repression (Lee and Schedl 2001; Jungkamp *et al.* 2011; Wright *et al.* 2011). We began by examining the expression of RME-2::GFP from *pwis116[rme-2p::rme-2::GFP::rme-2 3’UTR]* in oocytes at 22°C, to prevent transgene silencing. Again, the levels of RME-2::GFP were similar in the proximal oocytes of *sel-10(ar41)* and wild-type controls (Figure 7, G and H). Likewise, similar levels of endogenous RME-2 were seen in *sel-10(ok1632)* and wild-type oocytes stained with anti-RME-2-specific antibodies (Figure S12, I and J). Finally, we examined the expression of OMA-2, another well-established target of GLD-1-mediated translational repression (Lee and Schedl 2004; Wright *et al.* 2011; Scheckel *et al.* 2012). As we examined the expression of mNG::OMA-2 in *sel-10(ar41)* embryos in our analysis of LIN-41 Deg domains (Figure S7, A-D; described above), we quantitatively compared the expression level of mNG::OMA-2 expression in the proximal oocytes of the wild type and *sel-10(ar41)* mutants and observed a modest reduction (~30-50%) in mNG::OMA-2 expression levels in *sel-10(ar41)* mutants (Figure 7l). This result is agreement with the finding that an antibody that detects OMA-2 and its paralog OMA-1 exhibits a modest reduction in immuofluoresence staining (~10-33%, depending on the region of the proximal gonad analyzed) in the *sel-10(ok1632)* strong loss-of-function mutant (Kisielnicka *et al.* 2018).

Collectively, these results suggest that the ectopic GLD-1 in *sel-10* mutant oocytes is minimally effective at repressing translation of mRNA targets. The observation that some targets (e.g., *spn-4, mex-3,* and *rme-2)* might be unaffected by ectopic GLD-1, whereas others (e.g., *oma-2)* are modestly affected, is consistent with the observation that certain *gld-1* mutant alleles disrupt binding and repression of some mRNA targets but not others (Schumacher *et al.* 2005). Furthermore, these observations are again consistent with the fact that *sel-10* mutants are viable and fertile (Table 3), as the efficient repression of proteins such as SPN-4, MEX-3 and RME-2 during oogenesis should have negative consequences for embryonic development (Draper *et al.* 1996; Grant and Hirsh 1999; Gomes *et al.* 2001).

### The SCF^SEL-10^-dependent degradation of LIN-41 and GLD-1 depend on different kinases

As described above, the SCF^SEL-10^-dependent degradation of LIN-41 depends on CDK-1, but not MPK-1 (Figure S9, l-L). Consequently, we examined the requirement of these kinases for the SCF^SEL-10^-dependent degradation of GLD-1. Whereas *cdk-l(RNAi)* or *cdk-2(RNAi)* had no effect on the accumulation of GLD-1::GFP in proximal oocytes (n=14 and n=23, respectively), we observed ectopic expression of GLD-1::GFP in the proximal oocytes of *ozls5[gld-1::gfp]; mpk-l(ga111ts)* hermaphrodites at the non-permissive temperature (Figure S13). Thus, although both GLD-1 and LIN-41 are regulated by SCF^SEL-10^-dependent degradation, the temporal and spatial control of their accumulation during oogenesis is differentially responsive to protein kinase signaling, befitting their individual biological functions in promoting oogenesis.

## DISCUSSION

### Feedback regulation of LIN-41 and the spatial control of oocyte meiotic maturation

The oocytes of most sexually reproducing animals arrest in meiotic prophase for a prolonged period (reviewed by Huelgas-Morales and Greenstein 2017; Avilés-Pagán and Orr-Weaver 2018). This conserved arrest likely enables transcriptionally quiescent oocytes to grow by accumulating cellular organelles and cytoplasmic factors needed for embryonic development. Indeed, in *C. elegans* oocyte growth and meiotic maturation are coordinately controlled by LIN-41. In the absence of LIN-41 function, pachytene-stage oocytes abruptly cellularize, activate CDK-1, and enter M phase (Spike *et al.* 2014a). A salient feature of *C. elegans* oogenesis is that meiotic maturation is restricted to the oocyte in the most proximal position adjacent the spermatheca. This restriction ensures that only fully grown oocytes undergo meiotic maturation when they are positioned to enter the spermatheca during ovulation so they can become fertilized. Genetic evidence suggests that OMA proteins function to inhibit LIN-41 to facilitate meiotic maturation of the most proximal oocyte (Spike *et al.* 2014a). Specifically, proximal oocytes fail to enter M phase in *lin-41(ts); oma-l(null); oma-2(null)* triple mutants; whereas, pachytene stage oocytes prematurely enter M phase in *lin-41(null); oma-l(null); oma-2(null)* triple mutants (Spike *et al.* 2014a). Thus, the OMA proteins are absolutely required to spatially restrict for M-phase entry to the −1 oocyte, where they counteract LIN-41’s inhibitory activity. Consistent with this idea, molecular evidence suggests that LIN-41 is inactivated as a translational repressor in the final stages of oogenesis (Spike *et al.* 2014a; Tsukamoto *et al.* 2017), which precedes the elimination of LIN-41 upon the onset of meiotic maturation.

Specifically, two targets of LIN-41-mediated translational repression, *spn-4* and *meg-1,* are coexpressed with LIN-41 in the most proximal oocytes. The expression of *spn-4* and *meg-1* in proximal oocytes requires the function of the OMA proteins (Tsukamoto *et al.* 2017), consistent with the idea that the OMA proteins antagonize LIN-41 function in the late stages of oogenesis.

Here we show that the SCF^SEL-10^ promotes the rapid ubiquitin-mediated degradation of LIN-41 that leads to its elimination during meiosis I. Analysis of *sel-10* mutants indicates that the inactivation and degradation of LIN-41 are separable; the LIN-41 that accumulates in *sel-10* mutants appears to be largely inactive as a translational repressor. However, we did note that several LIN-41 variants (LIN-41(T83A) and LIN-41(ΔDeg-A)), which are defective in the SCF^SEL-10^-mediated degradation, decrease the meiotic maturation rate. This finding is consistent with the idea that LIN-41 inhibits meiotic maturation and that SCF^SEL-10^-mediated degradation constitutes a non-essential component of the regulatory mechanism. The nature of the “primary” mechanism inactivating LIN-41 prior to its degradation is currently unknown but could act on LIN-41 directly or another component of the large RNP complex it associates with (Spike *et al.* 2014b; Tsukamoto *et al.* 2017).

LIN-41 and CDK-1 reciprocally inhibit one another’s activity. Thus, the “primary” inactivation mechanism might play a key role in tipping the balance between LIN-41 and CDK-1 to generate a spatially restricted all-or-none meiotic maturation response. Upon its activation, CDK-1 triggers meiotic maturation and promotes the SCF^SEL-10^-dependent elimination of LIN-41. The elimination of LIN-41 requires the Deg-A and Deg-B domains in the LIN-41 N-terminal region. The LIN-41 Deg-A and Deg-B domains are intrinsically disordered and contain sequences that might function as phosphodegrons. The SCF^SEL-10^-dependent elimination of LIN-41 is blocked by the T83A mutation affecting a predicted CDK-1 phosphorylation site within the Deg-A domain, though whether this regulation is direct or indirect remains to be determined. We note one exception to the rule that CDK-1 activity promotes LIN-41 degradation. The *lin-41(tn1541tn1618)* mutation (Figure 2), which deletes the NHL domain, produces a strong loss-of-function *lin-41* mutant phenotype in which pachytene-stage oocytes enter M phase precociously. Nonetheless, we observe that the GFP::LIN-41(ΔNHL) protein still accumulates in the proximal gonad, albeit in an aberrantly punctate pattern (Figure S3, G-J). Interestingly, in the presence of a wild-type LIN-41 protein, the GFP::LIN-41(ΔNHL) mutant protein accumulates normally and is subject to SCF^SEL-10^-dependent degradation on schedule. It may be that the accumulation of the GFP::LIN-41(ΔNHL) protein in a punctate pattern correlates with its inaccessibility to CDK-l-dependent regulation. Alternatively, the degradation of LIN-41 during meiotic maturation may depend on LIN-41 activity during pachytene, as could be the case if a component of the SCF^sel-10^ degradation mechanism depends on *lin-41* function for its synthesis or activity.

The Deg domains may function as a timer to ensure that CDK-1 activity reaches an optimal threshold to ensure the successful completion of the meiotic divisions prior to the initiation of LIN-41 degradation. If LIN-41 is eliminated too early the fidelity of meiotic chromosome segregation may be compromised as is observed in certain hypomorphic *lin-41* mutant alleles (e.g., *tn1487tn1515, tn1487tn1516, tn1487tn1536,* and *tn1487tn1539;* Spike *et al.* 2014a). Thus, it will be important to elucidate the precise mechanisms by which the LIN-41 Deg domains link CDK-1 activity to SCF^SEL-10^-mediated degradation. The regulation of the Gl/S phase transition in budding yeast provides a framework for thinking about this issue (Nash *et al.* 2001; Kõvomägi *et al.* 2011; Yang *et al.* 2013; reviewed by Hopkins *et al.* 2017). The cyclin-dependent kinase complex, Cdk1-Clb5/6 promotes the entry into S phase but is inhibited by binding to its inhibitor Sic1 (Nugroho and Mendenhall 1994; Schwob *et al.* 1994). Sic1 is a substrate of the Cdk1-Clb5/6 kinase, which phosphorylates Sic1 to promote SCF^cdc4^-mediated degradation (Verma *et al.* 1997; Feldman *et al.* 1997; Nash *et al.* 2001). The cyclin-dependent kinase Cdk1-Clnl/2 initiates the decision to enter S phase during G1 and is not inhibited by Sic1. Phosphorylation of Sic1 by Cdk1-Clnl/2, while not sufficient to trigger Sic1 degradation, primes Sic1 for multisite phosphorylation by Clb5/6. The Sic1 CPD sequences contain multiple sites for phosphorylation by both Cdk1-Clb5/6 and Cdk1-Clnl/2, which results in the switchlike destruction of Sic1. A failure to degrade Sic1 substantially delays the Gl/S transition, whereas deletion of SIC1 causes DNA replication to initiate too early, resulting in genome instability (Nugroho *et al.* 1994; Cross *et al.* 2007). Further dissection of the mechanism by which the LIN-41 Deg domains function will illuminate whether analogous mechanisms are employed in a developmental context.

### Ubiquitin-mediated protein degradation and the OET

Signaling pathways and downstream kinase activation coordinate the cell-cycle and developmental events that underpin oocyte and early embryo development. In *C. elegans* the ERK MAP kinase signaling pathway and its effector kinase MPK-1 regulate pachytene progression and multiple aspects of oogenesis, including oocyte growth and specific events that occur during meiotic maturation (reviewed by Arur 2017). Consistent with these phenotypes, sustained activation of MPK-1 occurs during pachytene and in proximal oocytes (Lee *et al.* 2007). Likewise, in proximal oocytes activated cyclin-dependent kinase CDK-1 regulates an important aspect of oocyte meiotic maturation by promoting the transition from meiotic prophase to meiotic M phase, as we have described. Once activated, CDK-1 phosphorylates the DYRK mini-brain kinase MBK-2 as part of an intricate regulatory mechanism that permits MBK-2 activation near the end of the first meiotic division (Pellettieri *et al.* 2003; Stitzel *et al.* 2006, 2007; Cheng *et al.* 2009; Parry *et al.* 2009). These three kinases (MPK-1, CDK-1, and MBK-2) all function, at least in part, to promote the degradation of one or more RNA-binding proteins during oogenesis or the OET.

After meiosis, the OMA proteins are detectably phosphorylated by activated MBK-2 (Nishi and Lin 2005). Phosphorylation by MBK-2 promotes a direct physical interaction between the OMA proteins and the transcription factorTAF-4; this permits the sequestration of TAF-4 in the cytoplasm and prevents the premature onset of zygotic transcription (Guven-Ozkan *et al.* 2008). Furthermore, MBK-2-dependent phosphorylation primes the OMA proteins for phosphorylation by the glycogen synthase kinase GSK-3 and for degradation during the first mitotic division (Nishi and Lin, 2005; Shirayama *et al.* 2006). In addition to MBK-2 and GSK-3, the degradation of the OMA proteins requires the normal activities of additional kinases, including CDK-l/Cyclin B3, and several proposed E3 ubiquitin ligases (Shirayama *et al.* 2006; Du *et al.* 2015). The failure to degrade OMA-1 and eliminate it from early embryos is deleterious (Lin *et al.* 2003) and contributes to phenotypes exhibited by mutants that fail to degrade the OMA proteins (Shirayama *et al.* 2006). Indeed, the ectopic expression of OMA-1 in early embryos represses the translation of at least one mRNA target of the OMA proteins, *zif-1* mRNA, but only when OMA-1 is not phosphorylated by MBK-2, as in the *oma-l(zu405gf)* mutant (Guven-Ozkan *et al.* 2010). Thus, the MBK-2-dependent phosphorylation of the OMA proteins not only primes these proteins for degradation but also inhibits their ability to function astranslational repressors.

Likewise, the ability of GLD-1 to function as a translational repressor might be inhibited by MPK-l-dependent phosphorylation (Kisielnicka *et al.* 2018; this work). Since *mpk-1* activity is also required for the elimination of GLD-1, MPK-l-dependent phosphorylation would coordinate the inactivation of GLD-1 as a translational repressor with GLD-1 degradation. Consistent with this hypothesis, MPK-1 promotes the phosphorylation of GLD-1 and promotes its SCF^SEL-10^-mediated degradation (Kisielnicka *et al.* 2018; this work). Furthermore, this hypothesis potentially explains why the ectopic GLD-1 expressed in *sel-10* mutant oocytes is relatively ineffective at repressing the translation of multiple target mRNAs.

In sharp contrast to OMA-1 and GLD-1, our current understanding of the regulation of LIN-41 suggests that the inactivation of LIN-41 as a translational repressor is temporally and molecularly distinct from its degradation. Targets of LIN-41 translational repression such as *spn-4* and *meg-1* are actively translated prior to meiotic maturation and the CDK-l-dependent elimination of LIN-41. We have not yet determined that LIN-41 is phosphorylated by CDK-1 or any other kinase, as electrophoretic mobility changes are not reproducibly observed in *sel-10* mutants using several gel systems (unpublished results). The fact that *spn-4* and *meg-1* mRNAs are translated normally when *cdk-1* function is attenuated by RNAi (Tsukamoto *et al.* 2018) suggests that the CDK-1 is not required to inactivate LIN-41 as a translational repressor. In addition, we show here that mutations affecting the function of the LIN-41 Deg domains do not exhibit gain-of-function phenotypes or substantially repress the translation of LIN-41 target mRNAs.

### Multiple mechanisms regulate LIN-41 proteins

LIN-41 was first identified through its role in the heterochronic gene regulatory pathway that controls the timing of postembryonic cell divisions and cell fate decisions in somatic cells in *C. elegans* (Reinhart *et al. 2000;* Slack *et al. 2000;* reviewed by Rougvie and Moss 2013). In this capacity, LIN-41 functions to repress the translation of several transcription factors, including LIN-29, MAB-3, MAB-10, and DMD-3, which play key roles in specifying somatic cell fates during the L4 and adult stages (Reinhart *et al. 2000;* Harris and Horvitz 2011; Aeschimann *et al.* 2017). LIN-41 binds to the mRNAs of these genes and represses their translation during early larval stages (e.g., L1-L3) (Aeschimann *et al.* 2017). The Let-7 microRNA promotes the switch from early larval stages to the L4 and adult stages by repressing translation of LIN-41 beginning in the L4 stage (Reinhart *et al.* 2000; Slack *et al.* 2000). This regulation is specific to the soma as the *Iet-7(n2583ts)* mutation does not increase the accumulation of LIN-41 in the oogenic germline (Spike *et al.* 2014a). It is not clear whether specific protein degradation mechanisms collaborate with Let-7-mediated regulation to ensure that LIN-41 does not perdure from the early larval stages into the L4 and adult stage in somatic cells. If such mechanisms exist, they are unlikely to depend solely on the Deg domains because *lin-41* mutations affecting the Deg domains (e.g., *tn1620, tn1622 tn1635, tn1638, tn1643,* and *tn1645)* do not phenocopy *let-7* mutations or exhibit dominant somatic defects. Additionally, the Deg mutations do not confer an overt *lin-41(lf)* Dpy phenotype. Further, *Hn-41(tn1541tn1643[ΔDeg-A-RING-Deg-B])* L3-stage larvae do not exhibit precocious adult alae (n=7; Ann Rougvie, personal communication) as is frequently observed in *lin-41(n2914)* null mutants (Slack *et al.* 2000). The Deg domains mediate LIN-41 degradation during the OET over short time scales (i.e., 10-15 minutes), whereas the larval stages last for hours. This difference may obviate a requirement for SCF^SEL-10^-mediated degradation of LIN-41 during the larval stages. Interestingly, several *lin-41* gain-of-function alleles affecting the N-terminal 39 amino acid residues result in a defect in tip retraction during male tail development resulting in the production of a leptoderan (Lep) tail characteristic of other rhabditid nematode species (Del Rio-Albrechtsen *et al.* 2006). These *lin-41(Lep)* gain-of-function alleles do not affect LIN-41 degradation during the OET and thus define a site for LIN-41 regulation in somatic cells, which could involve proteolytic degradation.

LIN-41 is highly conserved. The mammalian ortholog LIN-41/TRIM71 is required for embryonic viability and neural tube closure in mice (Maller Schulman *et al.* 2008; Cuevas *et al.* 2015; Mitschka *et al.* 2015). LIN-41/TRIM71 was found to promote reprogramming of dermal fibroblasts to induced-pluripotent stem cells (IPSC) through the negative regulation of differentiation genes including the transcription factor EGR1 (Worringer *et al.* 2014). Importantly, the Let-7 microRNA inhibits reprogramming in part through the repression of LIN-41. Thus, the regulation of LIN-41 by Let-7 is a conserved regulatory module. By contrast, the Deg domains of *C. elegans* LIN-41 are not found in the mammalian orthologs and appear to be rapidly evolving in closely related rhabditid nematodes.

Developing systems must deploy mechanisms to extinguish RNA-binding protein-mediated translational repression. Such mechanisms may function to promote translation of batteries of genes needed to drive developmental transitions. LIN-41-associated mRNAs include many key genes required for embryonic development (Tsukamoto *et al.* 2017). Thus the inactivation of LIN-41 likely plays a key role in shaping the proteome during the OET. The “primary” mechanism inactivating LIN-41 prior to itsdegradation, and its potential conservation in LIN-41 orthologs or members of the TRIM-NHL class of RNA-binding proteins, remain to be determined.

## ACKNOWLEDGMENTS

We are grateful to Swathi Arur, Sarah Crittenden, Claire de la Cova, Daniel Dickinson, Bob Goldstein, Barth Grant, Iva Greenwald, Judith Kimble, Tim Schedl, and Dustin Updike for providing strains or reagents. We thank G. W. Gant Luxton for the use of his spinning disc confocal microscope. Some strains were provided by the Caenorhabditis Genetics Center, which is funded by grant P400D010440 from the NIH Office of Research Infrastructure Programs. We also thank WormBase for sequences and annotations. We thank Cynthia Kenyon for discussions on GLD-1 regulation. Ann Rougvie and Todd Starich provided helpful suggestions during the course of this work. This work was supported by NIH grant GM57173 (to D.G.).

